# Why variant effect predictors and multiplexed assays agree and disagree

**DOI:** 10.1101/2025.07.31.667868

**Authors:** Benjamin J. Livesey, Joseph A. Marsh

## Abstract

Computational variant effect predictors (VEPs) and multiplexed assays of variant effect (MAVEs) are two key tools for assessing the functional consequences of genetic variants. While their outputs are often concordant, there are also many differences. Here, we analyse missense MAVE data from 37 different human proteins, comparing them to five state-of-the-art VEPs in order to quantify and explain their points of agreement and disagreement. We find that discordance is not random but reflects fundamental differences in how each method infers functional impact. VEPs, which rely heavily on sequence conservation and basic structural features, tend to overcall pathogenicity at buried and hydrophobic residues, while underestimating impact in disordered regions and on charged surface residues. MAVEs, by contrast, capture context-specific mechanisms more accurately, but can miss pathogenic variants when the assay fails to reflect disease biology, or be subject to high levels of experimental noise. By comparing both global patterns and specific clinically relevant variants, we show how protein features, assay design, and variant type shape predictive discordance. Our findings provide a framework for interpreting when and why VEPs and MAVEs diverge and point toward strategies for improving variant interpretation through integrated, mechanism-aware approaches.

## Introduction

Genetic sequencing is becoming an ever more useful tool for medical diagnosis as advances have continued to make it more accessible and less expensive to perform. Unfortunately, our understanding of the mechanisms underlying the pathogenicity of the many novel variants being identified in disease-relevant proteins has not experienced the same amount of progress. This has led to a widening gap between the number of variants observed in sequencing data and the number with assigned clinical interpretations. As of December 2024, 57% of the single nucleotide variants (SNVs) reported in ClinVar (Landrum *et al*, 2018) that can be mapped to a UniProt primary isoform (The UniProt Consortium, 2025) protein sequence are of uncertain clinical significance, and that percentage is likely to rise as more sequencing is carried out. This also represents only a small fraction of SNVs possible in the sequence space of the human proteome.

Assays for protein variant function are a commonly employed tool to provide evidence for variant classification, however single-variant functional assays are too expensive and time-consuming to adequately fill the gap between the number of novel variants being observed and the need to identify those of clinical significance. To this end, advances have been made in the development and refinement of alternative high-throughput strategies to identify potentially pathogenic variants in the human population.

One such approach is the use of computational variant effect predictors (VEPs), also sometimes known as pathogenicity predictors. VEPs make use of various features such as sequence conservation, structural context and residue properties to predict the likelihood of a variant affecting protein function. While the methodology behind VEPs is diverse, they can be roughly divided into three classes based on their use of human population data. The largest class of VEPs is clinical-trained predictors. These VEPs include human population data in their models, often through training on labelled variants from databases such as ClinVar and HGMD (Stenson *et al*, 2020) but also potentially by exposure to human allele frequency data, often as a predictive feature. While clinical-trained predictors tend to perform well in benchmarks, they are also the most vulnerable to data circularity , and thus measurements of their performance may be inflated (Grimm *et al*, 2015).

Population-free VEPs have been steadily improving in performance over the past several years. These methods make no use of human population data either as training labels or features, often putting more emphasis on evolutionary conservation and structural features of proteins. Population-free predictors are not vulnerable to data circularity in their assessment and are competitive with clinical trained supervised predictors at the cutting edge (Meier *et al*, 2021; Laine *et al*, 2019; Frazer *et al*, 2021). The third category, population-tuned VEPs make some use of human population data for model tuning or optimisation, but do not use it as training labels or include allele frequencies as a feature. This is the smallest of the three classes and may be somewhat vulnerable to data circularity depending on the exact model training procedures, although the risk is smaller than with clinical-trained models. Compared to wet-lab functional characterisation, VEPs are highly cost effective (largely free-to-use) and produce results extremely rapidly, making them suitable for large-scale variant prioritisation. Determining the accuracy and reliability of VEP predictions is the subject of ongoing research like the Critical Assessment of Genome Interpretation (CAGI) (Critical Assessment of Genome Interpretation Consortium, 2024), and is highly dependent on the selected benchmark dataset (Livesey & Marsh, 2025).

Another approach is to use multiplexed assays of variant effect (MAVEs) (Fowler & Fields, 2014), a general term for a high-throughput assay in which the relative function of large numbers of genetic variants can be measured. MAVEs that focus on evaluating the relative fitness of protein coding missense variants may also be referred to as deep mutational scanning (DMS). In a typical MAVE for a protein coding gene, a library of variants is created in the gene of interest and then used to transform cells in which the function of the resultant mutant protein is linked to some measurable attribute such as cellular growth rate or reporter gene expression. The assay used to determine variant fitness can be fine-tuned to the specific protein or disease being assessed, for example, DNA binding activity for the PAX6 transcription factor (McDonnell *et al*, 2024) or protease activity of the caspase proteins (Roychowdhury & Romero, 2022). Many MAVEs use the relative growth rate of the variant-bearing cells as a proxy for fitness, if the function of the target gene is essential (Weile *et al*, 2017; Jiang, 2019; Sun *et al*, 2020). The relative fitness of each variant can be calibrated by using the measurements of null and wild type variants. Unlike VEPs, MAVEs represents a true measurement of protein activity or abundance in the cell following mutation, however it takes considerable expertise, time and specialist equipment to carry out, although it is considerably less time consuming and expensive than performing separate functional assays for each individual variant. Several MAVEs have reported extremely good performance for the classification of variants from ClinVar (Findlay *et al*, 2018; Grønbæk-Thygesen *et al*, 2024; Gersing *et al*, 2023).

Both MAVEs and VEPs are different tools that can be used to address the same issue: the annotation of variants of uncertain clinical significance (VUS). Indeed, in the American College of Medical Genetics and Association for Molecular Pathology (ACMG/AMP) guidelines for variant interpretation, the two different annotation sources can be combined with one another to improve evidence strength (Richards *et al*, 2015). Currently VEPs are only capable of providing supporting evidence for pathogenicity while functional assays such as MAVEs can provide up to strong evidence. Significant advances in both fields have been made since the 2015 guidelines were published, and work is under way to produce a set of guidelines with notably improved evidence strengths for VEPs (Pejaver *et al*, 2022; Badonyi & Marsh, 2025). With the understanding that evidence for the reclassification of many VUS are likely to come from both MAVE and VEP sources in the near future, it is important to know how the two approaches differ for the purpose of combining evidence from them. Can we really treat functional measurements from a MAVE and functional predictions from a VEP as independent of each other if they are both attempting to estimate the same value?

We have recently shown that VEPs, in particular recently developed population-free methods show a strong correlation with MAVE datasets across numerous proteins (Livesey & Marsh, 2025). We used this relationship as the basis of three VEP benchmarking studies, highlighting the progress in VEP development across several years. The best-performing predictors approached a Spearman’s correlation of 0.8 with some MAVE datasets. While both approaches are attempting to address the same fundamental questions of variant effects, the extremely different methodologies employed are naturally going to result in contradicting predictions in some proteins and protein regions. It is important both for future computational method development and integration of the two methods as separate lines of evidence to know where and why the two approaches disagree on the effects of variants on protein function.

In this study we compare and contrast the performances of 37 MAVE datasets and 5 representative VEPs to determine the extent and causes of discordance between the predictions made by the different approaches. We identify protein regions in which MAVEs and VEPs tend to differ in their predictions and general properties that may be predictive of such regions in other proteins. Finally, we assess several cases of known pathogenic and benign variants where MAVEs and VEPs produce contradicting results and deduce the reasons for the discrepancies.

## Results and discussion

### MAVE and VEP dataset selection and normalisation

The MAVE datasets used in this study are the same 36 MAVE datasets we previously used to benchmark 96 VEPs (Livesey & Marsh, 2025) with the addition of a *BRCA1* dataset (Findlay *et al*, 2018), which was excluded from the benchmark due to the large overlap between scored variants and variant databases (Table 1). The mutagenesis technique, fitness assay, cellular model system and general technique employed by the MAVE experiments is diverse, although yeast-based functional complementation assays are one of the most common (Weile *et al*, 2017; Gersing *et al*, 2023; Sun *et al*, 2020). Other notable experimental setups include VAMP-seq assays for protein stability (Clausen *et al*, 2024; Matreyek *et al*, 2018), a saturation genome editing approach in a human cell line (Findlay *et al*, 2018), binding assays (Bandaru *et al*, 2017; Weng *et al*, 2024) and several unique assays tailored for protein-specific functions (Miller *et al*, 2022; Amorosi *et al*, 2021). These datasets were curated to ensure that the fitness assay employed is relevant to normal protein function or stability. A single score-set is selected to be representative of each MAVE target protein, even in cases where multiple groups produced results for a single protein. The selected score set is the one with the highest median correlation to 96 VEPs in the benchmark study.

**Table 1.** Summary of 37 MAVE datasets.

| Gene/UniProt | Cellular Model | Expression strategy | Fitness assay | Total variants * | Pathogenic/Likely pathogenic ** | Benign/Likely benign ** | Reference |
| --- | --- | --- | --- | --- | --- | --- | --- |
| ADRB2 / P07550 | Human HEK293-derived cell line | Plasmid integration | Reporter gene expression | 7800 | 0 | 4 | (Jones *et al*, 2020) |
| ASPA / P45381 | Human HEK293 cells | Plasmid integration | VAMP-seq | 5843 | 50 | 6 | (Grønbæk-Thygesen *et al*, 2024) |
| BRCA1 / P38398 | Human HAP1 cells | Saturation genome editing | Relative growth rate | 1837 | 168 | 77 | (Findlay *et al*, 2018) |
| CALM1 / P0DP23 | *S. cerevisiae* | Plasmid expression | Relative growth rate | 1813 | 16 | 0 | (Weile *et al*, 2017) |
| CARD11 / Q9BXL7 | Human lymphoma cell line | Saturation genome editing | Relative growth rate | 2395 | 7 | 1 | (Meitlis *et al*, 2020) |
| CASP3 / P42574 | *E. coli* | Plasmid expression | Activity assay (GFP signal) | 1507 | 0 | 1 | (Roychowdhury & Romero, 2022) |
| CASP7 / P55210 | *E. coli* | Plasmid expression | Activity assay (GFP signal) | 1680 | 0 | 2 | (Roychowdhury & Romero, 2022) |
| CBS / P35520 | *S. cerevisiae* | Plasmid expression | Relative growth rate | 10450 | 105 | 4 | (Sun *et al*, 2020) |
| CYP2C9 / P11712 | humanized yeast strain YMD4256 | Plasmid expression | Click-seq (activity assay) | 6142 | 0 | 1 | (Amorosi *et al*, 2021) |
| GCH1 / P30793 | *S. cerevisiae* | Plasmid expression | Relative growth rate | 3482 | 13 | 1 | Not yet published |
| GCK / P35557 | *S. cerevisiae* | Plasmid expression | Relative growth rate | 8570 | 260 | 7 | (Gersing *et al*, 2023) |
| GDI1 / P31150 | *S. cerevisiae* | Plasmid expression | Relative growth rate | 7589 | 3 | 2 | (Silverstein *et al*, 2022) |
| HMBS / P08397 | *S. cerevisiae* | Plasmid expression | Relative growth rate | 5963 | 39 | 2 | (van Loggerenberg *et al*, 2023) |
| HMGCR / P04035 | *S. cerevisiae* | Plasmid expression | Relative growth rate | 16853 | 1 | 0 | (Jiang, 2019) |
| HRAS / P01112 | E. coli | Plasmid expression | Binding assay | 3135 | 26 | 4 | (Bandaru *et al*, 2017) |
| KRAS / P01116 | S. cerevisiae | Plasmid expression | Binding assay | 1185 | 36 | 0 | (Weng *et al*, 2024) |
| LDLRAP1 / Q5SW96 | S. cerevisiae | Plasmid expression | Binding assay | 5795 | 0 | 3 | (Jiang, 2019) |
| MAPK1 / P28482 | Human A375 cells | Viral integration | Relative growth rate | 6810 | 11 | 0 | (Brenan *et al*, 2016) |
| MSH2 / P43246 | Human HAP1 cells | Viral integration | Relative growth rate | 16749 | 126 | 238 | (Jia *et al*, 2021) |
| MTHFR / P42898 | S. cerevisiae | Plasmid expression | Relative growth rate | 12445 | 31 | 9 | (Weile *et al*, 2021) |
| NUDT15 / Q9NV35 | Human HEK293T cells | Plasmid integration | Relative growth rate | 2934 | 0 | 0 | (Suiter *et al*, 2020) |
| PAX6 / P26367 | S. cerevisiae | Plasmid expression | (Transcription factor) activity assay | 2709 | 36 | 1 | (McDonnell *et al*, 2024) |
| PDE3A / Q14432 | Human GB1 cells | Viral integration | Relative growth rate | 7893 | 1 | 3 | (Garvie *et al*, 2021) |
| PPARG / P37231 | Human THP-1 cells | Viral integration | Downstream gene expression | 9595 | 13 | 3 | (Majithia *et al*, 2016) |
| PPM1D / O15297 | Human K562 cells | Viral integration | Downstream gene expression | 7889 | 1 | 14 | (Miller *et al*, 2022) |
| PRKN / O60260 | Human HEK293T cells | Plasmid integration | VAMP-seq | 8756 | 14 | 15 | (Clausen *et al*, 2024) |
| PTEN / P60484 | S. cerevisiae | Plasmid expression | Activity assay | 6564 | 188 | 4 | (Mighell *et al*, 2018) |
| SERPINE1 / P05121 | E. coli | Plasmid expression | Phage display/binding assay | 1988 | 0 | 2 | (Huttinger *et al*, 2021) |
| SHOC2 / Q9UQ13 | Human cell line | Viral integration | Relative growth rate | 10972 | 3 | 6 | (Kwon *et al*, 2022) |
| SNCA / P37840 | S. cerevisiae | Plasmid expression | Relative growth rate | 1787 | 3 | 2 | (Newberry *et al*, 2020) |
| SRC / P12931 | S. cerevisiae | Plasmid expression | Relative growth rate | 3372 | 0 | 0 | (Ahler *et al*, 2019) |
| SUMO1 / P63165 | S. cerevisiae | Plasmid expression | Relative growth rate | 1700 | 0 | 0 | (Weile *et al*, 2017) |
| TP53 / P04637 | Human H1299 cells | Viral integration | Relative growth rate | 2967 | 171 | 30 | (Kotler *et al*, 2018) |
| TPK1 / Q9H3S4 | S. cerevisiae | Plasmid expression | Relative growth rate | 3181 | 4 | 1 | (Weile *et al*, 2017) |
| TPMT / P51580 | Human HEK293T cells | Plasmid integration | VAMP-seq | 3689 | 0 | 1 | (Matreyek *et al*, 2018) |
| UBE2I / P63297 | S. cerevisiae | Plasmid expression | Relative growth rate | 2563 | 0 | 0 | (Weile *et al*, 2017) |
| VKORC1 / Q9BQB6 | Human HEK293T cells | Plasmid integration | VAMP-seq | 2695 | 1 | 2 | (Chiasson *et al*) |

\* The total number of single amino acid substitution variants in the protein covered by the MAVE assay.
\*\* The number of ClinVar variants with a 1\* or higher review status covered by the MAVE assay.

Compared to modern VEPs; older, well-known VEPs such as SIFT and PolyPhen-2 are largely redundant, so we selected five state-of-the-art VEPs to perform our analysis. We previously found each of these methods to be either immune or highly resistant to the phenomenon of data circularity, so making comparisons using annotated disease variants should not materially affect the results. AlphaMissense is an advanced language model built on AlphaFold2 that makes use of protein structure and is tuned with human population frequencies (Cheng *et al*, 2023). ESCOTT is a combination of the conservation-based statistical GEMME model (Laine *et al*, 2019) and further structural features (Tekpinar *et al*, 2024). ESM-1v is a pure language model designed specifically for zero-shot variant effect prediction (i.e. no protein-specific training) (Meier *et al*, 2021). SaProt is a model that combines the 3D structural context of residues with their sequence context, producing a hybrid sequence/structure language model (Su *et al*, 2024). VARITY_R is a top-performing clinical-trained predictor that is relatively insensitive to data circularity and sources training data from several databases (Wu *et al*, 2021). Together, these five predictors represent a diverse methodological cross-section of VEP techniques at the cutting edge of current methodology. They are also all either population-free predictors or, in the case of VARITY and AlphaMissense, not particularly sensitive to data circularity. This is important as data circularity may influence VEP outputs based on previously encountered clinical data, potentially skewing comparisons with MAVE data, especially in the context of previously observed clinically relevant variants.

MAVE and VEP performance is heterogenous across proteins. To demonstrate this, we calculated the area under the receiver operating characteristic curve (AUROC) for each of the five selected predictors in addition to the MAVE data. We used ClinVar pathogenic variants with a 1-star or higher review status as the pathogenic dataset and gnomAD as the putatively benign dataset. While gnomAD is not a purely benign set of variants, we refer to it as ‘putatively benign’ as it provides a representation of genetic variation in a healthy human population. While gnomAD certainly contains recessive and low-penetrance pathogenic variants, particularly among variants with low allele frequency, it better reflects the real-life clinical application of VEPs than other purely benign alternatives, that being distinguishing rare pathogenic variants from rare benign variants. Datasets like ClinVar-benign, as well as containing far fewer variants, are primarily composed of common variants in the human population, as allele frequency is a major determining factor in the benign classification criteria. The AUROC calculation was bootstrapped 1000 times by sampling with replacement from both datasets independently to obtain a distribution (Figure 1). Depending on the protein and/or MAVE assay, the MAVE data can either perform worse than the VEPs (*HMBS*, *GCK*, *MTHFR*), similarly to the VEPs (*GCH1*, *PAX6*, *PPARG*) or better (*MSH2*, *TP53*, *BRCA1*) in separating the pathogenic variants from the putatively benign. The five VEPs selected for this analysis have similar levels of performance, and their outputs also strongly correlate with one another. They have a much lower, although still significant, correlation to the MAVE assay data (Figure S1). This demonstrates that VEPs are much more similar to each other than to the MAVEs, despite their varying methodologies and are likely to identify similar types of pathogenic variants to one another.

**Figure 1.**
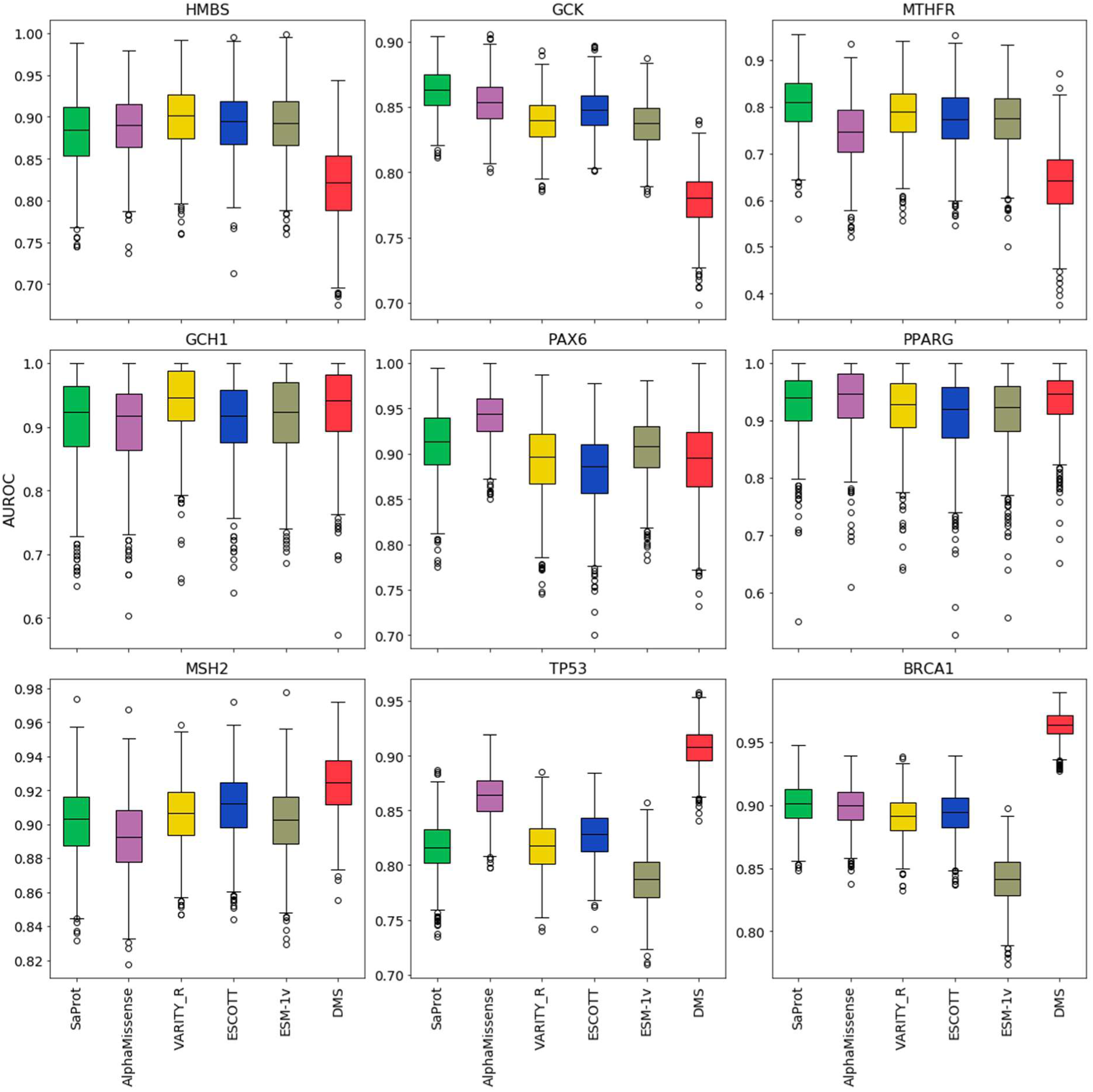
The area under the receiver operating characteristic curve (AUROC) for the five VEPs and MAVE dataset across nine proteins. The pathogenic dataset was comprised of ClinVar pathogenic and likely pathogenic variants with a review status of 1 star or higher. The putatively benign dataset was composed of gnomAD variants not present in the pathogenic set. Only proteins with at least 10 variants from each dataset were considered. The distributions plotted were produced by bootstrapping the AUROC calculation 1000 times, sampling with replacement independently from both datasets. The same variants were used to calculate the AUROC for each VEP and the MAVE data on each bootstrap.

The first challenge in characterising areas of proteins where VEPs and MAVEs disagree is making the various scoring systems employed comparable. Most commonly, VEPs assign scores on a 0-1 scale, where the number indicates the probability of a mutation being damaging to protein function, however this is far from the only scoring system. MAVE datasets most frequently use a relative fitness scale with wild-type-like variants scoring 1 and null variants 0, which is the opposite of the most common VEP scale but also allows for worse-than-null and fitter-than-wild type variants unlike most VEPs. Comparing scores that use such a diverse range of scales is challenging, which is why we decided to use normalised rank scores.

To perform normalised ranking, we took each score scale and first ensured that lower values represent the fitter variants by deducting every score from 1 if the scale was inverted. We then ordered the scores by value and assigned each variant a rank on a per-protein basis. The ranks were then normalised between 0 and 1, providing a simple and comparable scale for all datasets. In cases where multiple scores are being compared and some are missing data, the missing data is removed from all score sets before applying the normalisation, ensuring that rankings are always performed with the same variants in each score set.

A comparison can be made between any two MAVE or VEP score sets for a particular protein by deducting one set of normalised rank scores from the other, resulting in values close to 0 where the two sets agree perfectly and values up to 1 or -1 in the case of extreme discordance between the scores. Regions of discordance as measured by the difference between normalised rank scores can then be associated with protein structural features, sites and domains, providing insight into the features of proteins that MAVEs and VEPs are most likely to disagree on. This approach is illustrated in Figure 2, where predictions from the ESM-1v VEP are compared to a set of deep mutational scan results for the low density lipoprotein receptor adaptor protein 1 (Jiang, 2019), and plotted alongside important protein features.

**Figure 2.**
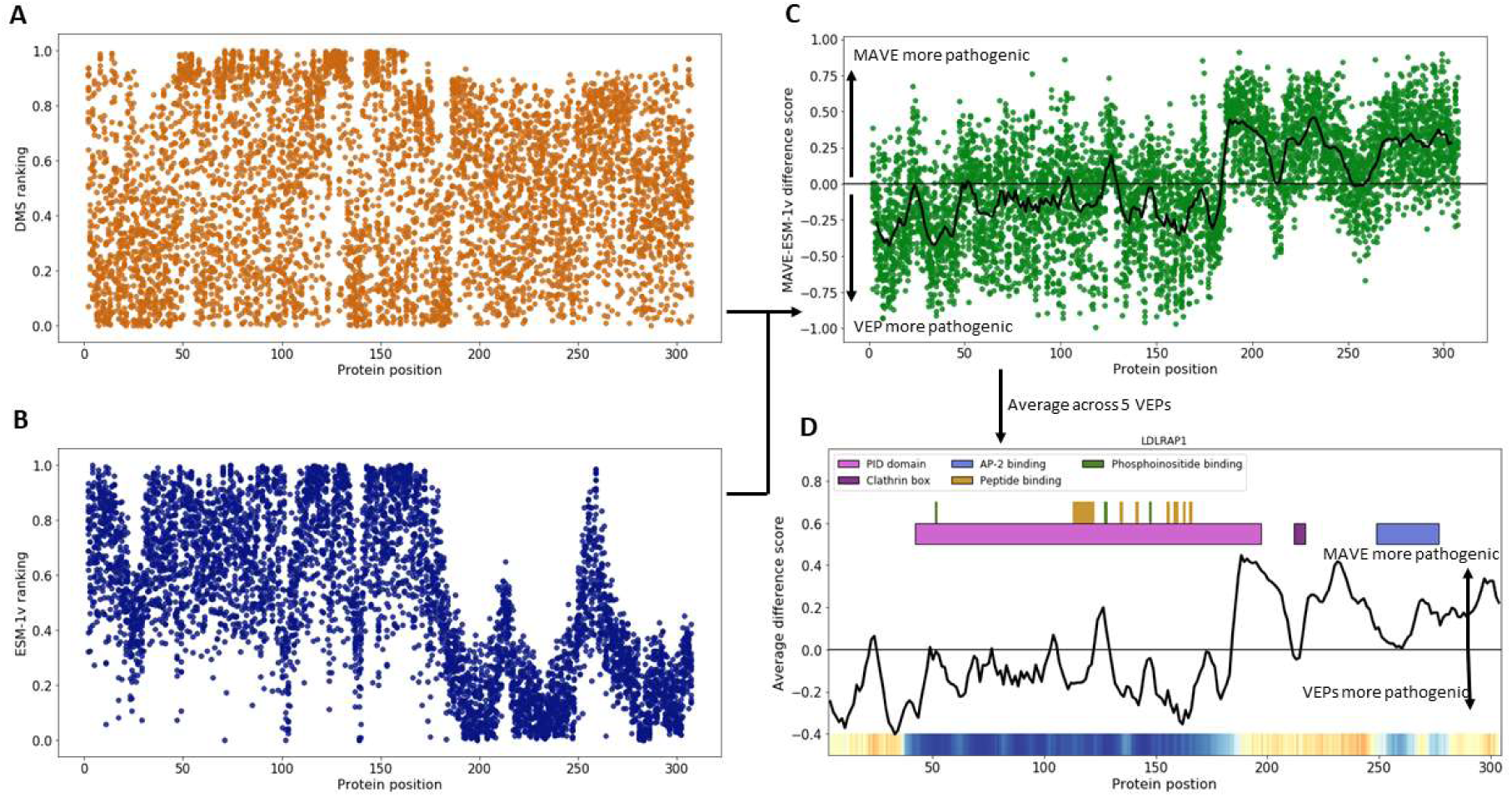
An illustration of how comparisons are made between VEP and MAVE datasets on different scales. **A**. The MAVE dataset for the protein LDLRAP1 is converted into ranks and normalised between 0 and 1. **B**. The predictions from ESM-1v for LDLRAP1 are also converted into ranks and normalised between 0 and 1. **C.** The data in section (B) is deducted from the data in section (A) to give the difference in normalised ranks across the protein. The black line represents a moving average across the protein for a window of 100 variants (∼5 residues) with a stride size of 25 variants. Higher scores represent variants that the MAVE finds to be more impactful on protein function than the MAVE, while lower scores indicate the opposite. **D**. The mean of the normalised difference, averaged across all five VEPs (black line) is compared to the location of protein features (top) and disordered regions as indicated by AlphaFold pLDDT (bottom).

The first notable difference between MAVE and VEP datasets can be observed in the distribution of normalised rank scores. As illustrated in Figure 2A and 2B, scores for different amino acid substitutions at each protein position in the MAVE data tend to be dispersed across a wider range of values than for the ESM-1v predictor. This pattern remains consistent across other VEPs and MAVE datasets (Figure S2) with normalised MAVE rank scores showing greater standard deviation at each residue position and also across small windows of 10 amino acids than the VEPs. VARITY was excluded from this figure, as it only produces predictions for variants possible by single nucleotide substitutions (∼30% of all variants) and thus naturally demonstrates lower positional deviation than the other VEPs. Interestingly one dataset stood out as having particularly low standard deviation per-position, the MAVE set for *PPARG* (Majithia *et al*, 2016), with an average standard deviation of only 0.0763, compared to the MAVE average of 0.196 (excluding *PPARG*).

### Insight into amino acid level predictions

It is well known that mutations involving certain amino acids are much more likely to be damaging to protein function than others (Petukh *et al*, 2015). For example, aromatic amino acids are very bulky and hydrophobic, often driving the formation of the hydrophobic core of globular proteins. Mutations of these large residues to those with different properties, or inclusion of these residues on the protein surface can lead to altered folding pathways or misfolding (Chiti & Dobson, 2006). Cysteine residues are vital in the formation of disulphide bridges, particularly in extracellular proteins and those localised to oxidising compartments such as the endoplasmic reticulum. Additionally, the unique conformational constraints of proline and the flexibility of glycine mean that substitution at these positions often lead to significant changes in protein backbone structure.

We investigated the differences between MAVEs and VEPs in terms of their tendency to label mutations involving specific amino acids as more or less damaging to protein function. To perform this analysis, we pooled data from the five representative VEPs and compared them to the MAVE data. Our results indicate that there are some significant differences between the tendency of VEPs and MAVEs to identify specific amino acid changes as damaging.

We present the Hedges G effect size of the differences in the normalised rank scores for the pooled MAVE data against the pooled VEP data for every combination of wild type and mutant amino acid in Figure 3 as a heatmap. Positive (red) effect sizes indicate variants that the MAVE tends to label as more damaging than the VEP while negative (blue) effect sizes indicate variants that the VEPs identify as more damaging. The intensity of the colour indicates the magnitude of the effect.

**Figure 3.**
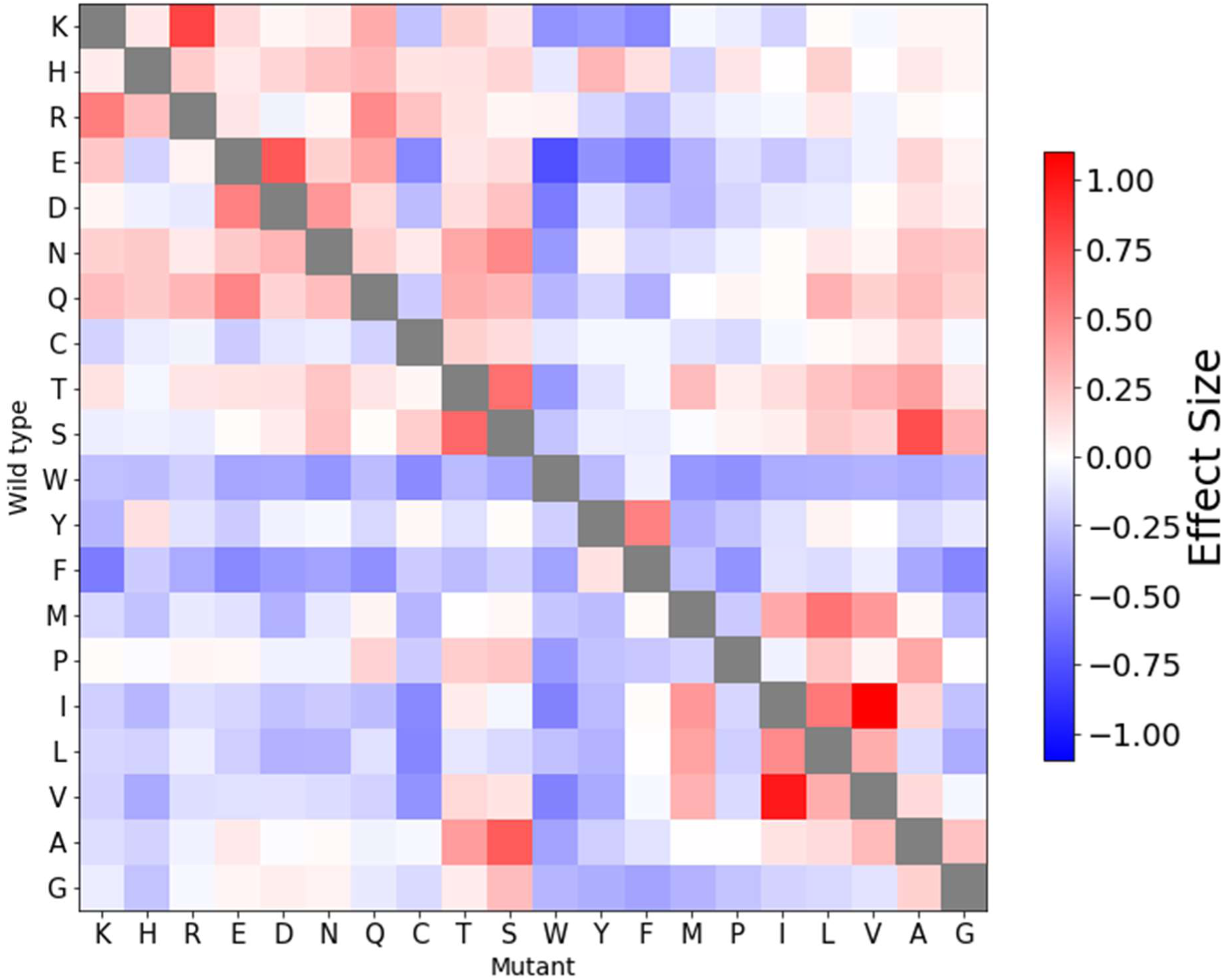
A heatmap of the effect sizes between normalised VEP and MAVE ranks for all amino acid substitutions. Positive, red effect sizes indicate substitutions that MAVEs fund more pathogenic than the VEPs while negative, blue effect sizes indicate substitutions that VEPs find more pathogenic than the MAVEs. White squares indicate substitutions where both methods are concordant. Synonymous mutations are not included in this analysis and the grey boxes indicate blank squares.

There are several interesting trends that can be observed on this heatmap. First, the plot is generally mirrored along the diagonal, indicating a consistent magnitude and direction of the effect size regardless of the orientation of the wild type and mutant residue. I.E., an isoleucine to valine mutation and a valine to isoleucine mutation are both predicted more damaging by MAVEs than VEPs by about the same amount on average.

The upper left quadrant of the heatmap, representing substitutions between charged and polar residues shows almost all positive effect sizes (except cysteine). This indicates that MAVEs tends to find these substitutions between residues with somewhat similar properties to be more damaging than VEPs do. Similarly, there is also a cluster of small, hydrophobic residues on the bottom right (I, L, V, A) where substitutions between them also demonstrate a similar positive effect size and thus a tendency to be labelled as more damaging by MAVE data than by VEPs.

Almost any mutation involving the bulky, hydrophobic amino acids (W, Y, F) as either the wild type or the mutant results in negative effect sizes and are thus found more damaging by the VEPs than by the MAVE. A similar effect can also be observed in the bottom left quadrant, representing variants where small, hydrophobic residues are mutated to charged or polar ones.

The overall pattern is fairly clear: mutations that are considered to be *conventionally* damaging to protein function, i.e., hydrophobic to charged, small to bulky or bulky to small are routinely predicted as more damaging by the VEPs than MAVEs. Mutations that are *conventionally* considered as less likely to be disease-causing, i.e., charged to charged or small/hydrophobic to small/hydrophobic are found to be more damaging by the MAVEs than the VEPs. This general pattern is consistent across different subsets of the top-performing VEPs and MAVE datasets, demonstrating that the trend is not being driven by a small set of MAVE datasets or VEPs.

Our speculation as to the reason for this pattern is as follows. MAVEs make no prior assumption about the effects of amino acid substitutions but bases the outcome purely on an independent fitness assay that is unique for each protein and fitness definition. VEPs have to make assumptions (during their training process) about the types of variants that would typically be damaging in order to be generalisable across many different proteins. At the core of most VEPs is a multiple sequence alignment, providing residue-level conservation features to the predictor, that will generally reflect known evolutionary trends of bulky and property-altering substitutions being less favourable than those between residues with similar properties. By learning these trends, VEPs become better able to generalise their predictions across multiple diverse proteins but may be unable to identify occasional exceptions to these trends in different protein contexts that MAVEs are able to identify consistently. In other words, VEPs are likely overcalling substitutions that are typically considered the most damaging and undercalling those that are most often phenotypically neutral while MAVEs are producing a more balanced overview.

This also ties in with our observation that MAVEs tends to assign a greater range of scores to variants at a single position in a protein than VEPs do. The prediction made by VEPs is based on their previous experience (training), which includes the fact that some wild type amino acids are less tolerant of certain mutations than others in general. MAVEs are unique for each protein, leading to a larger range of predictions at each position that is unconstrained by prior assumptions of residue-level mutational tolerance.

### Protein features with discordant VEP/MAVE predictions

In each of the 37 MAVE datasets, we identified the subset of variants with the largest gap between the five representative VEPs and MAVEs in terms of normalised rank differences. We term the 5% of variants with the largest normalised rank difference where the VEP assigns the variants as more damaging than the MAVEs as VEP-damaging variants and those the MAVE assigns as more damaging as MAVE-damaging variants. We also identified the 5% of variants with the smallest absolute normalised rank difference, representing those where both MAVE and VEPs concur about the relative functional impact of the variants (concordant variants).

#### VEP-damaging variants are enriched among buried residues and interface residues

We used a simple, previously described method to classify residues as belonging to the protein surface, interior or interface and break down interface residues into core, support or rim (Levy, 2010) using biological assemblies in the PDB (Berman *et al*, 2000). We found previously that interface rim residues resembled protein surface residues more than the interface core or support in terms of properties and pathogenic mutation enrichment (Livesey & Marsh, 2022), so we grouped them with the protein surface residues. VEP-damaging variants occur most frequently in protein interior and interface regions, while MAVE-damaging variants are depleted (Figure 4A). The opposite pattern is observed in protein surface residues where VEP-damaging variants are depleted while MAVE-damaging variants are enriched. This finding also ties in with our previous observation that the substitution of bulky hydrophobic residues, which are most commonly found in the hydrophobic core of globular proteins, are often predicted as more damaging by VEPs than MAVEs. Charged and polar residues are more common on the protein surface than the interior, and mutations of these are frequently predicted as more damaging by MAVEs than VEPs.

**Figure 4.**
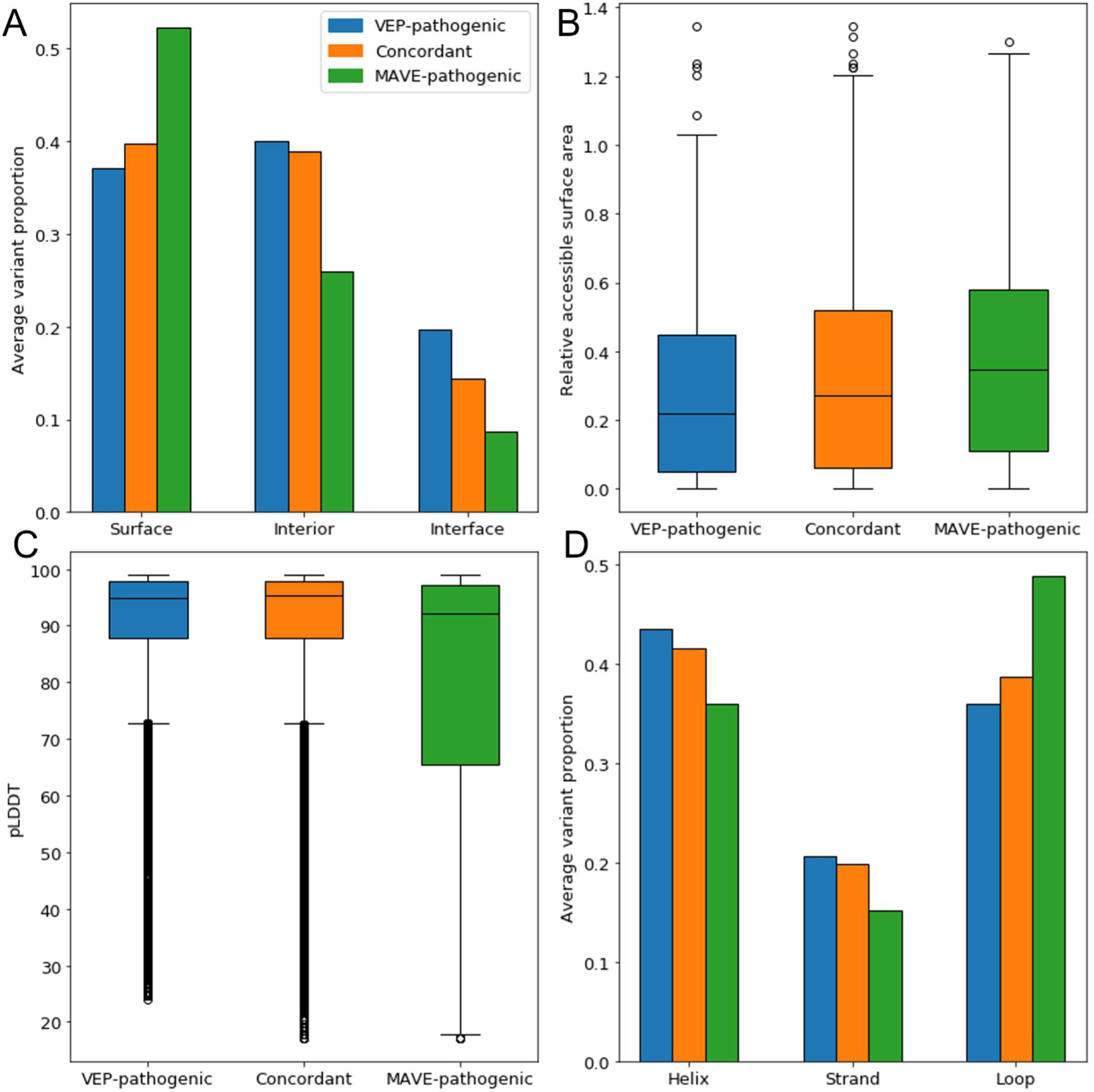
Characteristics of discordantly predicted mutations. **A.** The proportion of variants from the VEP-pathogenic, concordant and MAVE-pathogenic classes occurring at the protein surface, interior and interface residues (interface rim residues are included in the surface group). **B.** The distribution of relevant solvent accessibility within the three classes. **C**. The distributions of AlphaFold-derived predicted local distance difference test (pLDDT) values within the three classes. **D.** The proportion of variants from each class that that occur within helices, strands and loops as defined by DSSP.

We also calculated the relative solvent accessibility (RSA) of each residue using AlphaFold-predicted structures of every protein and compared the RSA of the three classes (Figure 4B). Residues where VEP-damaging mutations occur have a lower RSA than the other two classes, indicating that this class contains the highest proportion of buried residues. MAVE-damaging mutations have the highest RSA and are thus likely primarily surface residues. Those variants where MAVEs and VEPs are concordant are intermediate to the other two classes, likely indicating a mix of interior and surface residues.

#### MAVE-damaging mutations are enriched in disordered regions

We obtained predicted local distance difference test (pLDDT) values from AlphaFold2-generated PDB files. pLDDT is a per-residue measurement of confidence in the AlphaFold2 structural prediction, and regions of low pLDDT correspond to disordered regions of proteins. While the median pLDDT for all three groups of variants is above 90, meaning that the residue positions were confidently predicted on average, the MAVE-damaging group has both the lowest median (92.27) and a distribution where the third quartile extends to much lower pLDDT values than the other two groups (Figure 4C). It has been observed that VEPs tend to produce poorer predictions of pathogenicity within intrinsically disordered regions of proteins (Fawzy & Marsh, 2025). Disordered regions are usually less well conserved throughout evolution than structured protein regions. Evolutionary conservation is also a vital component of almost all VEPs, and sometimes the only feature employed (particularly for ESM-1v), under the assumption that variants in poorly conserved regions are more likely to be benign.

While it is true that disordered regions of proteins are highly enriched in benign variants relative to pathogenic ones, VEPs tend to excessively classify the majority of variants in disordered regions as benign. MAVEs are under no such constraints in disordered regions, so it is likely that MAVE data are closer to the biological reality of variant effects in disordered regions. The observed trend is being driven by VEPs preferentially predicting the majority of variants in disordered regions as benign, resulting in variants found even moderately damaging by the MAVEs being discordantly predicted and appearing in the MAVE-damaging variants.

We also used the Define Secondary Structure of Proteins (DSSP) algorithm to classify residues as belonging to a helix, a strand or a loop (no defined secondary structure) (Figure 4D). VEP-damaging variants were enriched relative to MAVE-damaging variants in both helices and strands, while MAVE-damaging variants were enriched in loops. This pattern is also likely related to protein disorder, as most disordered regions will be included within the loop class, while helices and sheets are unlikely to harbour disordered regions.

### Characteristics of discordantly predicted protein regions

In order to determine the protein regions that corresponded to the greatest levels of discordance between the VEPs and the MAVEs, we extracted protein feature annotations from UniProt including notable domains, motifs and protein-protein interaction sites, we also used InterPro to annotate all conserved regions and residues from the Conserved Domain Database (CDD). We plotted the mean normalised difference between the MAVE and VEPs across the protein sequence and structure in order to determine the protein features with the greatest discordance between predictions of pathogenicity (Figure 5 and Supplemental file SF1). Also shown are the pLDDT scores per residue obtained from AlphaFold, which is a useful proxy for sequence disorder.

**Figure 5.**
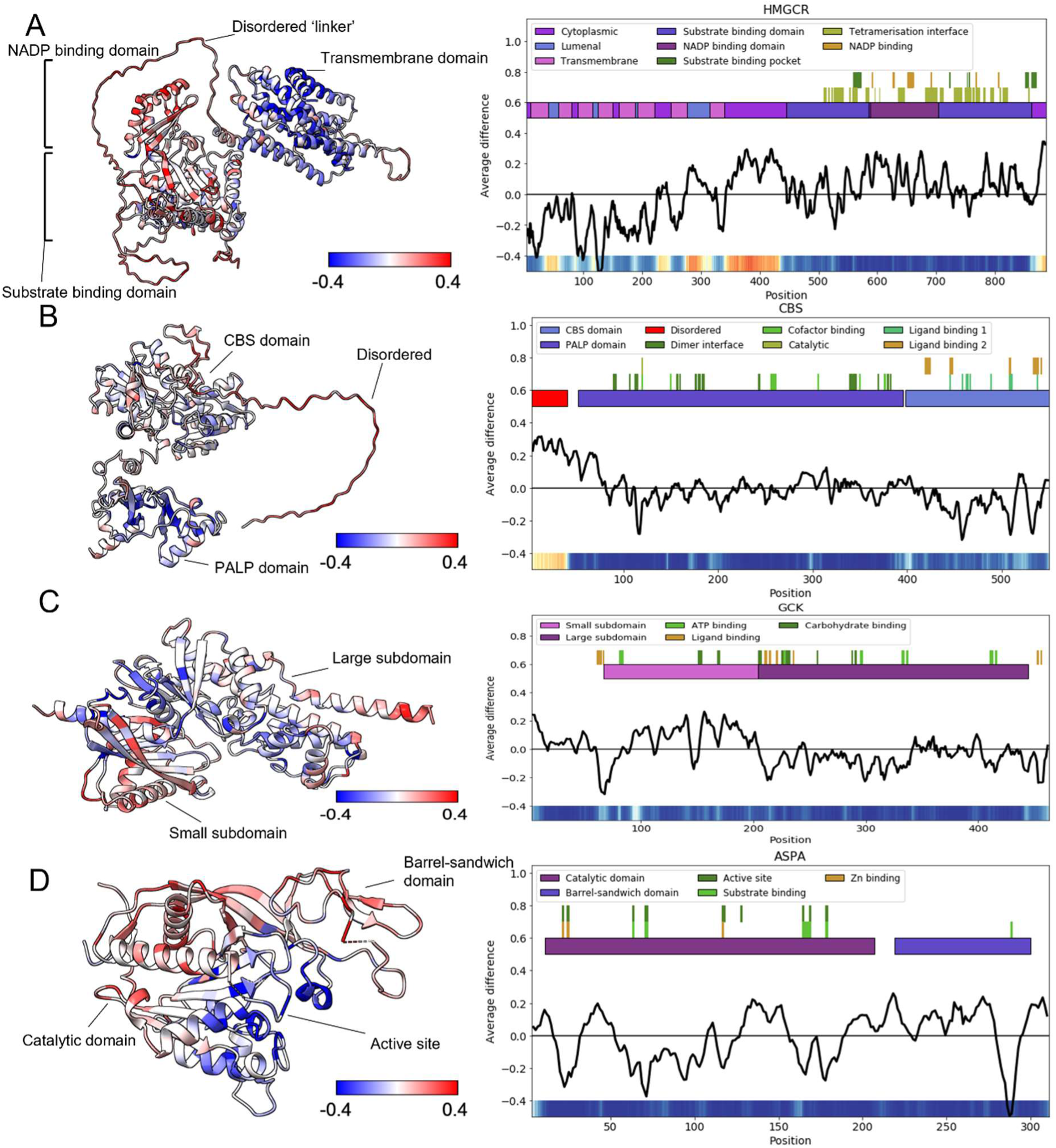
The correspondence of normalised rank difference to protein sequence and structural features. Each panel displays the Alphafold2-derived structure of the protein with residues coloured by the mean normalised difference between VEPs and MAVEs. Higher (red) values indicate residues where the MAVE considers variants more pathogenic than the VEPs, while lower (blue) values indicate residues where the VEP predicts variants to be more pathogenic than the MAVE. Alongside is a plot, where the rectangles at the top represent protein domains, regions and conserved residues. The black like represents the mean normalised difference between VEPs and DMS within a scanning window covering 100 datapoints (at least 5 residues). The heatmap at the bottom of each plot represents the AlphaFold2 predicted local distance difference test (pLDDT) scores across the protein with blue representing greater confidence in the structure prediction, and red representing lower confidence. **A.** Structure and plot for *HMGCR*. **B.** Structure and plot for *CBS* **C.** Structure and plot for *GCK*. **D.** Structure and plot for *ASPA*. Structures and plots for the remaining 33 proteins are available in supplemental file S1.

Within all residues covered by each extracted annotation, we calculated the overall average of the mean normalised difference value across all possible variants between the MAVE and the VEPs. Thus, regions with a negative average difference value are predicted to be more pathogenic by the VEPs, and regions with a positive average difference value are found to be more pathogenic by the MAVE. The full results of this analysis are available in Table S1.

In several cases, entire protein domains were preferentially predicted as pathogenic by either the MAVE or the VEPs, demonstrating a systematic, domain wide difference between the predictions and measurements made by the two methods. This was most noticeable in the sterol-sensing domain of HMGCR (residues 61-218) which has an average difference of -0.22, indicating that the VEPs identify variants in this region as significantly more pathogenic than the MAVE (Figure 5A). A similar effect was notable in the CBS domain of CBS (418-476) where the average difference was - 0.12 (Figure 5B). There are also several domains that exhibit a positive average difference, albeit to a lesser extent including the PPARG nuclear receptor ligand binding domain (238-503) with an average difference of 0.07 and the GCK hexokinase small subdomain (67-203) with an average difference of 0.08 (Figure 5C).

Of the twelve proteins in our dataset with Uniprot-annotated disordered regions present, all had positive average differences within those regions ranging from 0.03 (*GCH1*) to 0.22 (*CBS,* Figure 5B), indicating that variants in these regions are consistently found more pathogenic by the MAVEs than by the VEPs. Disordered regions of proteins typically have low sequence constraint and harbour notably fewer damaging variants and more functionally neutral ones than ordered regions (Ahmed *et al*, 2022). These two attributes mean that VEPs tend to perform poorly at identifying the few truly damaging variants present within these regions (Fawzy & Marsh, 2025) as sequence conservation is a core component of almost every VEP (and all of those used for this analysis). Furthermore, the high frequency of neutral variants may also lead to supervised VEPs that have been trained using variants in these regions learning to associate disordered regions with benign protein variation. It is therefore likely that this pattern observed in disordered regions is being driven by poor VEP performance in those regions and a preference to label most variants as benign. MAVEs performance is independent of evolutionary conservation, and thus MAVEs are likely providing more reliable measurements of variant function in regions of disorder.

Transmembrane helices are annotated in three of the proteins in our dataset (*HMGCR*, *VKORC1* and *ADRB2*). All transmembrane regions (with the sole exception of *ADRB2* transmembrane helix 4) have a negative difference value, indicating that variants in these regions are predicted as more pathogenic by VEPs than by the MAVEs (Figure 5A). Transmembrane helices span the hydrophobic interior of a lipid bilayer and are thus predominantly composed of hydrophobic residues, even more so than helices on the protein interior (Hildebrand *et al*, 2004). While not classified as such in our protein interface data, transmembrane helices can be thought of as large protein-ligand interfaces. The similar amino acid composition of transmembrane regions to the protein interior is likely resulting in the VEPs treating these features like part of the protein interior where variants that significantly alter properties of hydrophobic residues may have a deleterious impact on protein folding. While variants in transmembrane helices have the potential to impact membrane integration, they are far less likely to affect the normal protein folding pathways than variants in helices within the protein interior and are thus likely less damaging than predicted by the VEPs. The MAVEs likely represent a more balanced overview of the functional impact of variants in transmembrane domains.

Conserved active site residues are among the most concordantly predicted residues among all of the MAVE datasets. 15 proteins have annotated active sites within the UniProt and InterPro annotations, and of these, the 9 have average difference values between 0.07 and -0.06 (*HMGCR*, *SRC*, *NUDT15*, *MAPK1*, *PDE3A*, *PTEN*, *GCH1*, *CASP3* and *CASP7*). This makes sense as active site residues tend to be highly conserved, thus variants would impact both MAVE measurements of activity and VEP predictions. 5 other proteins (*ASPA*, *PRKN*, *VKORC1, TPK1* and *MTHFR*) all have low negative average difference values at the annotated active sites. For three of these proteins, the enrichment is easily explained by looking at the fitness assay employed during the MAVE assay. *PRKN*, *ASPA* and *VKORC1* were all assayed used VAMP-seq, a method that assesses relative fitness by quantifying protein expression (identifying structurally disruptive mutations) and does not directly quantify protein activity (Matreyek *et al*, 2018). Thus, catalytically impaired variants will not be identified by the MAVE assay unless they also cause the protein to misfold. This mismatch between the MAVE readout and the fitness predictions being made by the VEPs demonstrates that caution should be used when employing MAVE as a true fitness standard as the definition of protein fitness in the assay may not match precisely with mechanisms of pathogenicity in the protein. An extreme case of this is shown in Figure 5D for *ASPA* where variants in residues within and in close proximity to active site are all predicted more pathogenic by the VEPs than by the DMS. In *TPK1* and *MTHFR*, the reasons for the negative average difference at active site residues is less clear.

Large average difference values are also notable among dimerisation, protein interaction, metal and ligand binding sites (Table S1). Although the magnitude and direction of the difference varies from protein to protein.

### Analysis of discordantly predicted clinical variants

While it is clear that there are systematic differences between MAVE and VEP measurements of variant effect, it is less clear which approach is producing the more reliable results without referring to a ‘gold standard’. For this purpose, we created two sets of labelled human variants for the 37 proteins.

- The Clinvar-pathogenic dataset is composed of human single nucleotide polymorphisms annotated as Pathogenic or Likely pathogenic in the ClinVar database. We retained only records with a 1-star review status or higher.
- The Clinvar-benign dataset is constructed similarly to the Clinvar-pathogenic one, but including only variants annotated as Benign and Likely benign.

We pooled the normalised rank differences across each of the MAVE datasets. We investigated variants from the clinvar-pathogenic and clinvar-benign datasets with extremely high or low (>0.75 or <-0.75) mean normalised rank differences, being those with the greatest level of discordance between VEPs and MAVEs and attempt to identify a possible cause for the disagreement with respect to its pathogenic/benign label. Figure 6 shows the normalised MAVE scores plotted against the mean of the normalised VEP scores for nine proteins with discordantly predicted variants. We have performed a detailed analysis of the variants in the top left and bottom right of this plot, being those that are most discordantly predicted by MAVEs and VEPs. The normalised rank differences for all Clinvar-pathogenic and Clinvar-benign variants across the 37 MAVE datasets is available in Table S2.

**Figure 6.**
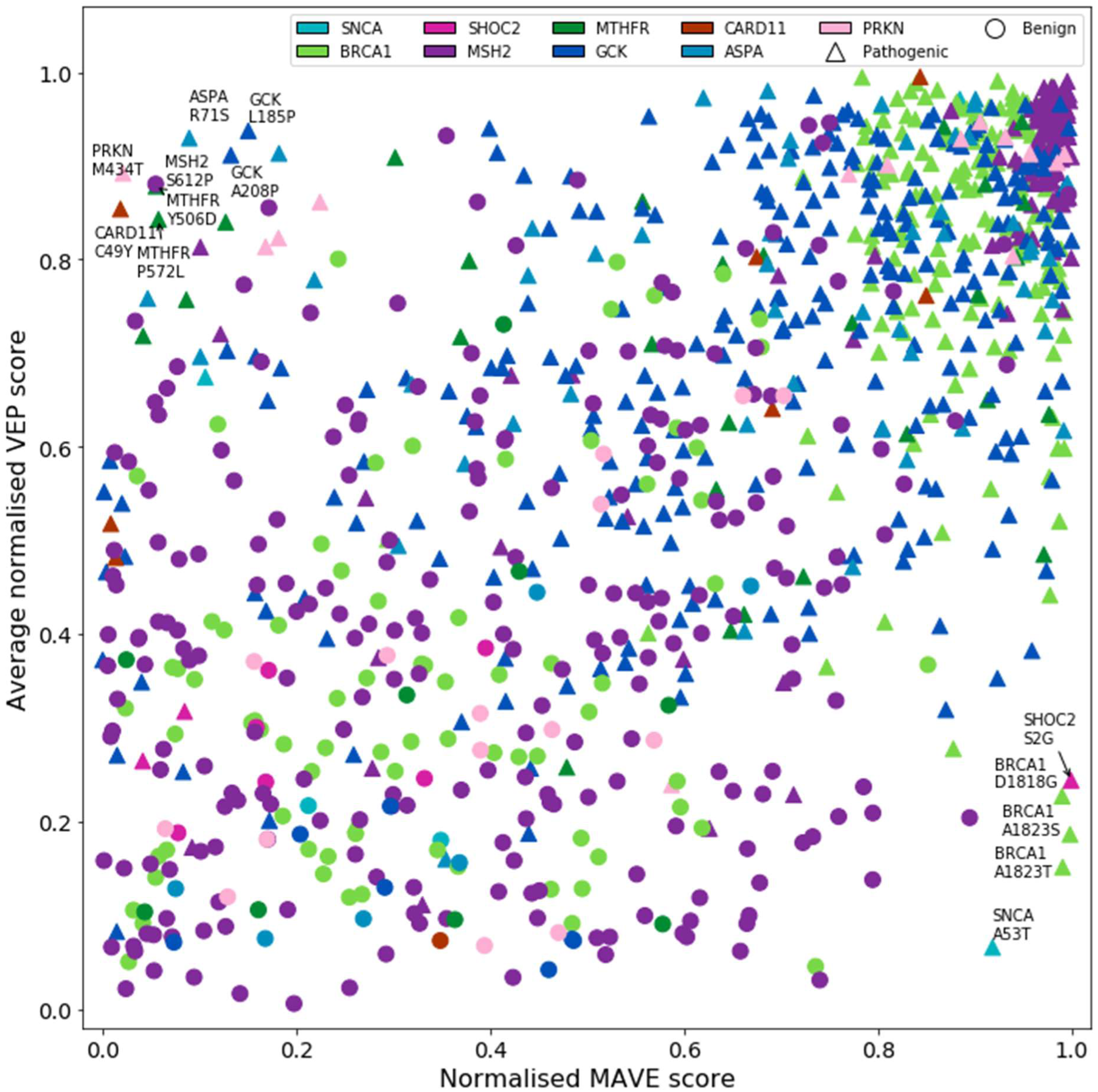
Normalised MAVE score plotted against the mean of the normalised VEP scores for ClinVar-pathogenic and ClinVar benign variants in nine proteins. The labelled variants are those that are most discordantly predicted as measured by the mean normalised difference between MAVEs and VEPs.

#### SNCA

α-synuclein is a protein that plays an important role in synaptic activity. Variants in α-synuclein can cause Parkinson’s disease and other neural degenerative conditions collectively called α-synucleinopathies (Alafuzoff & Hartikainen, 2017). Pathogenic variants typically trigger aberrant aggregation of α-synuclein into structures such as Lewy bodies that impair normal cellular functions. The MAVE for α-synuclein activity was performed in yeast cells with the readout being the yeast growth rate, which is impaired by, and thus a proxy for, α-synuclein aggregation (Newberry *et al*, 2020).

While there is only a total of five ClinVar variants in α-synuclein after filtering for entry quality, it contains the ClinVar-pathogenic variant with the highest mean normalised rank difference (0.853), A53T. This is a variant found pathogenic by the MAVE but predicted as benign by all VEPs. The A53T mutation is well known to have a high aggregation potential and is implicated in Parkinson’s disease (Lee *et al*, 2002). Using an alignment of 1051 sequences related to α-synuclein generated by MMSeqs2 (Steinegger & Söding, 2017), we found that threonine is actually the most common residue at the equivalent position to alanine 53 in the human sequence (401 occurrences vs only 84 for alanine) (Figure S3A). The most likely explanation is that VEPs are failing to correctly predict this mutation because the pathogenic mutant residue is common at this position through evolution.

Given that this a well-known pathogenic mutation with extensive characterisation, the VEPs are unquestionably incorrect in their assessment of variant effect, likely because this mutation violates their baseline assumption that evolutionary conservation is a good proxy for variant effect. It may be that this variant is only pathogenic in the context of humans due to our relatively long lifespan, given that Parkinson’s disease has a late age-of-onset, typically appearing after the age of 60.

#### BRCA1

Breast cancer type 1 susceptibility protein performs a central role in the homology-directed repair of DNA double-strand breaks (Moynahan *et al*, 1999). In addition, *BRCA1* also participates in non-homologous end-joining DNA repair (Baldeyron *et al*, 2002) and performs several other functions relevant to maintaining genome stability, it is classed as a tumour suppressor. Pathogenic variants in *BRCA1* increase the risk of breast and/or ovarian cancer, typically through loss of protein function.

The MAVE for *BRCA1* used a saturation genome editing approach to introduce variants into the protein’s N-terminal RING domain and C-terminal BRCT domains in the HAP1 human cell line (Findlay *et al*, 2018). *BRCA1* is an essential gene in HAP1 cells, and loss of function impairs cellular growth, permitting a readout of protein function by measuring cellular growth rate.

Of the 245 ClinVar variants in *BRCA1*, only three exceed the 0.75 threshold: A1823T, A1823S and D1818G, all of which are pathogenic. The MAVE concurs with the ClinVar classification and also finds all three pathogenic, while the VEPs assign all three as benign. The codon encoding A1823 straddles two exons, so the variants at that location may impact splicing. Pangolin (Zeng & Li, 2022) predicts a 65% chance of splice donor loss on a threonine mutation (NM_007294.4(BRCA1):c.5467G>A (p.Ala1823Thr)) and a 67% chance of splice loss on a serine mutation (NM_007294.4(BRCA1):c.5467G>T (p.Ala1823Ser)). A study has found that D1818G also impacts splicing and results in skipping of exon 22 (Rouleau *et al*, 2010, 22). It is likely the VEPs are failing to account for the impact on splice sites, resulting in incorrect benign predictions for these variants.

Because this MAVE was performed using saturation genome editing in a human cell line, the protein was able to be studied in its endogenous context, with native splicing machinery. A similar assay performed in yeast, or even using expression from a plasmid in human cell lines would almost certainly be insensitive to such splice-impacting variants, similarly to the VEPs.

#### SHOC2

Leucine-rich repeat protein SHOC-2 is a scaffold protein and a core component of the SHOC2-MRAS-PP1c (SMP) complex that regulates MAP kinase signalling. Pathogenic mutations in *SHOC2* are known to be a cause of several RASopathies, most notably Noonan syndrome-like disorder with loose anagen hair 1 (Garavelli *et al*, 2015). This condition is typified by characteristic, slow growing, sparce and easily pluckable hair in addition to the developmental, cognitive and cardiac defects that characterise RASopathies. The MAVE for *SHOC2* was performed in a KRAS-mutant human cancer cell line by introducing the mutant sequence with a viral vector and knocking out the native SHOC2 gene (Kwon *et al*, 2022). Fitness was determined by the relative abundance of each variant at the endpoint of the assay.

*SHOC2* has nine variants in ClinVar, which largely show good concordance with VEP predictions of pathogenicity. The one exception to this is S2G which is a ClinVar pathogenic mutation, found pathogenic by the MAVE, but predicted benign by the VEPs. This particular variant is known to introduce an N-myristoylation site into the protein, resulting in aberrant targeting to the plasma membrane (Cordeddu *et al*, 2009). With the exception of Serine at position 2, the N-terminus of *SHOC2* satisfies the consensus sequence for N-myristoyltransferase targeting. As part of its normal function, SHOC2 is translocated from the cytoplasm into the nucleus under growth factor stimulation, but is unable to effectively do so when targeted to the plasma membrane by N-myrisoylation, explaining the poor variant fitness in the MAVE. It is also unsurprising that VEPs failed to identify this pathogenic gain-of-function mutant given the small number of proteins to which such a site can be introduced by a single missense variant.

#### MSH2

The primary role of DNA mismatch repair protein Msh2 is to locate and repair DNA mismatches, which is a function it performs as a heterodimer with MSH6, forming the MutSα complex. Alternatively, MSH2 can dimerise with MSH3, forming the MutSβ complex which repairs larger-scale insertions and deletions (Traver *et al*, 2015). Pathogenic variants in MSH2 cause Lynch syndrome, an autosomal dominant disease that greatly increases cancer susceptibility. Many of the variants that cause Lynch syndrome result in decreased mismatch repair activity (Ollila *et al*, 2006). The MAVE for MSH2 was carried out in human HAP1 cells, with endogenous expression of MSH2 knocked out, mutant MSH2 was introduced using a lentiviral construct (Jia *et al*, 2021). Selection was performed using a purine analogue that is toxic to cells with a functional mismatch repair pathway, resulting in higher growth rates for cells with reduced MSH2 activity (I.E., a reverse-survival assay, where cells with reduced protein fitness are better able to proliferate than the wild type).

A single variant in mismatch repair protein *MSH2* exceeds the concordance threshold, S612P. This is a likely benign variant in ClinVar which is identified as benign by the MAVE, but as pathogenic by the VEPs. This variant occurs in an alpha helix, which proline mutations tend to disrupt, and in an alignment of 13624 related sequences, only five occurrences of proline were observed at this position, demonstrating strong residue constraint (Figure S3B). We suggest that the MAVE may actually be incorrect and that this may actually be a mis-labelled variant in ClinVar because, in addition to the likely helix disruption and constraint at this residue, no individuals bearing this variant are present in gnomAD and numerous other variants in the same helix are ClinVar-pathogenic (A604D, D603Y/G/V, A600D/V). A closer look at the category assignment shows that the *MSH2* MAVE study is actually used as functional evidence supporting the classification of likely benign (VCV002447337.2 - ClinVar - NCBI), further complicating the comparison and raising the question of data circularity when using the ClinVar classification as the gold-standard in this case.

#### MTHFR

Methylenetetrahydrofolate reductase is a metabolic enzyme that catalyses the conversion of 5,10-methylenetetrahydofolate to 5-methyltetrahydrofolate, which serves as a methyl group donor for the conversion of homocysteine into methionine (Froese *et al*, 2018). Variants that cause a severe loss of MTHFR function (usually defined as <20% of wild type function remaining) cause homocystinuria, an autosomal recessive disease that results in developmental delay and severe intellectual disability (Burda *et al*, 2015). The MAVE for *MTHFR* makes use of the ability of the protein to functionally complement its yeast homologue *MET13,* which is essential for yeast growth unless supplemental methionine is added to the growth medium (Weile *et al*, 2021). Variant function was measured by assessing the yeast growth rate via a Tile-seq approach.

Y506D and P572L are likely pathogenic and pathogenic/likely pathogenic variants in ClinVar respectively, found benign by the MAVE and predicted as pathogenic by the VEPs. Y506 is a core residue at a dimerisation interface. This particular variant was not remarked on by the authors of the MAVE study but it was noted that dimerisation interface residues scored no lower than other residues at the surface of the same domain (Weile *et al*, 2021). This was attributed to dimerisation not being strictly required for *MTHFR* activity, although the pathogenic classification of the variant implies that some potentially unaccounted-for function of the protein is impacted by mutation of the interface. One patient study found that a compound heterozygote bearing the Y506D variant along with a partial duplication in the other allele (p.Leu89_Pro101dup) had only 8.1% *MTHFR* activity remaining (Burda *et al*, 2015). P572 is a fully buried residue, it is reported as pathogenic and likely pathogenic in ClinVar by two reporters, and additionally has been found to severely reduce MTHFR activity in a human expression system (3.1 ±0.4% activity remaining) (Burda *et al*, 2017). It is likely that, in both cases that the VEPs have successfully identified a pathogenic variant that was overlooked by the MAVE, possibly due to experimental noise, or a difference between the yeast expression system and human gene context.

#### GCK

Hexokinase-4 is a metabolic enzyme that phosphorylates hexose sugars at the first stage of glycolysis (Takeda *et al*, 1993). It also acts as a glucose sensor in pancreatic beta cells. Pathogenic variants in GCK are associated with defects in glucose metabolism, specifically maturity-onset diabetes of the young 2 (MODY2) which is an early-onset autosomal dominant disease where insulin secretion is impaired. The MAVE for GCK activity was carried out using yeast complementation in a yeast strain in which the three yeast hexokinase genes had been deleted (Gersing *et al*, 2023). This strain was unable to grow on glucose medium, but was rescued by the activity of plasmid-expressed human GCK.

There are two likely pathogenic variants in GCK that are predicted pathogenic by the VEPs but found to be benign by the MAVE: A208P (reviewed by expert panel) and L185P. Neither variant is present in gnomAD, and all other variants at L185 are found by the MAVE to result in severe loss-of-function. Most other variants as A208 also result in a loss-of-function. It is highly likely that the benign MAVE results are due to experimental noise in this case and are not accurately reflecting protein function.

#### CARD11

Caspase recruitment domain-containing protein 11 is a signalling protein involved in the adaptive immune response following the activation of T and B cell surface receptors. It is activated by phosphorylation, homooligomerises and recruits *BCL10* and *MALT1*, leading to the phosphorylation of downstream proteins and eventual release of NF-kappa-B (Gaide *et al*, 2001). *CARD11* variants can cause B-cell expansion with NFKB and T-cell anergy, an autosomal dominant condition characterised by splenomegaly (enlarged spleen) and polyclonal B-cell expansion (Snow *et al*, 2012). *CARD11* variants can also cause both dominant and recessive forms of immunodeficiency (Greil *et al*, 2013; Ma *et al*, 2017). The MAVE for *CARD11* activity used a saturation genome editing approach in a human B cell lymphoma cell line, TMD8, where growth is dependent on NF-κB signalling (and therefore upstream *CARD11* and B-cell receptor signalling as well) (Meitlis *et al*, 2020). As *CARD11* activity is vital to growth in this cell line, relative fitness was determined by the abundance of each variant following selection.

C49Y is a pathogenic mutation in ClinVar that is found benign by the MAVE and predicted as pathogenic by the VEPs. C49 is a hotspot for gain-of-function variants in *CARD11*, where multiple variants increase downstream NF-κB activity (Meitlis *et al*, 2020). The difference between MAVE and VEP predictions at this location is caused by a disconnect in readouts between the two approaches. VEPs are (accurately) predicting the variant as pathogenic, while the MAVE is (accurately) capturing the increased downstream activity caused by the variant – however this fitter-than-wild type phenotype just looks benign in a direct comparison of the scores due to the rank normalisation. In this particular fitness assay, gain-of-function variant do not negatively impact cellular fitness as they would in the context of the whole organism. A solution that can help flag pathogenic gain-of-function variants is to use an approach employed by Weile et al. and ‘flip’ these scores by setting them to their reciprocal score (assuming a 1-0 scale), we have found that this approach improves MAVE correlation with VEPs in a previous study (Livesey & Marsh, 2020).

#### ASPA and PRKN

Aspartoacyclase is an enzyme that catalyses the deacetylation of N-acetylaspartic acid to form acetate and L-aspartate, it is thought to regulate the concentration of N-acetylaspartic acid in brain tissue (Herga *et al*, 2006). Pathogenic variants, resulting in severe loss of enzyme activity cause Canavan disease, a rare lethal neurodegenerative condition that often results in death in early childhood (Mendes *et al*, 2017).

E3 ubiquitin ligase parkin functions as part of an E3 ubiquitin ligase complex, and catalyses the covalent attachment of ubiquitin proteins onto its targets (Shimura *et al*, 2000). PRKN plays a role in the degradation of misfolded proteins by attaching chains of ubiquitin, targeting them to aggresomes. Variants in this protein that impair its ligase activity can lead to both to classic Parkinson disease and an autosomal recessive form of Parkinson disease (Hedrich *et al*, 2002, 50; West *et al*, 2002).

The MAVEs for both ASPA and PRKN were performed using the VAMP-seq technique which assesses variant abundance through quantified expression of a GFP fusion protein (Grønbæk-Thygesen *et al*, 2024; Clausen *et al*, 2024). Unlike most other MAVE assays discussed here, VAMP-seq does not measure protein activity, but rather protein stability since unstable variants will be rapidly degraded and therefore not contribute to the GFP signal produced by the cell.

R71S and M434T in *ASPA* and *PRKN* respectively are both ClinVar pathogenic mutations found benign in the MAVEs and predicted as pathogenic by VEPs. Because VAMP-seq was used, stability altering mutations were identified, but not activity-altering mutations that did not also disrupt protein stability, which is likely the case for these two variants. The mutation in *ASPA* alters a residue that binds the proteins substrate, likely abrogating the binding (Bitto *et al*, 2007). The mutation in *PRKN* occurs in close proximity to the active site and may also affect substrate binding.

It is clear that VEPs can fail in situations where their assumption of evolutionary conservation being tied to phenotypic impact is violated. This could occur in proteins that have developed relatively new functions compared to their homologues throughout evolution or in ‘new’ and thus poorly evolutionarily conserved areas of proteins. Similar effects are likely in protein sequences that are under strong positive selection as in numerous immune related proteins. VEPs are known to best predict loss-of-function variants (Gerasimavicius *et al*, 2022), which makes their failure to identify the N-terminal gain-of-function mutation in *SHOC2* and the splicing variants in *BRCA1* understandable.

MAVE data are subject to experimental noise, some of which will be cleaned up during replication or various statistical filtering procedures, but at least some low-quality results will inevitably make it into the final datasets. While it is difficult to positively confirm whether or not experimental noise is the cause of any given case of discordance with VEP predictions, it seems likely that many of the cases where the MAVE results diverge from the ClinVar label and the VEP accurately predicts the label may be due to noise in the MAVE results. The other major cause of MAVE prediction failure is when the assessed protein function or readout from the MAVE does not match the mechanism(s) underlying the disease being investigated. We see this in both *ASPA* and *PRKN*, that the VAMP-seq assay for protein stability fails to identify variants that reduce protein activity but maintain stability. This downside can be ameliorated by performing multimodal assays instead of studying variant effects in a single context, using a single assay to produces fitness scores. Multimodal assays can be tailored to the unique mechanisms underlying each disease related to a protein, and have the potential to provide significant insight into the pathogenicity of variants that standard MAVE assays may fail to identify.

## Conclusion

Understanding both the strengths and limitations of variant interpretation methods is essential for improving clinical classification of variants of uncertain significance. Our analysis highlights key systematic differences between VEPs and MAVEs. VEPs tend to overcall pathogenicity in structurally buried regions, including protein interiors, interfaces, and transmembrane domains, while undercalling potentially pathogenic variants on protein surfaces or within intrinsically disordered regions. MAVEs, in contrast, can provide direct functional evidence but are limited by experimental noise and the choice of assay, which may not capture all disease-relevant mechanisms.

Expanding the use of MAVEs to assay distinct phenotypes is critical. A single assay may miss important variant effects. For example, VAMP-seq may overlook catalytic impairment at active sites. Applying different assay modalities allows protein functions to be dissected more precisely, offering insights into gain-of-function, dominant-negative, or splicing mechanisms that would otherwise go undetected. Saturation genome editing (SGE) represents a major step forward in this area, enabling variant assessment within native genomic contexts and avoiding artefacts introduced by overexpression systems.

With the continued improvement of computational variant effect predictor technology, correct prediction of cases similar to those highlighted here where the VEP failed to correctly identify a ClinVar labelled variant will become more important. A remarkable amount of progress has been made by focussing purely on evolutionary conservation, and VEPs that solely make use of this feature remain among the top performers to date (although in the case of *SNCA,* the over-reliance on conservation also proved a hinderance). Several recent predictors have benefitted from the addition of protein structural features on top of conservation (Gerasimavicius *et al*, 2025), implying that the path to further progress may be from integration of alternative sources of informative features. One such feature that may lead to improvements is the addition of a splice-site impact predictor, which would have allowed the VEPs to correctly identify the three splice-impacting variants in BRCA1 as pathogenic.

From the analysis of mutation level preferences, and the performance of VEPs in both disordered regions and transmembrane helices, it does appear that MAVEs tend to be the more reliable methods in general. Most of the quirks of VEP predictions we identified are rooted in their requirement to generalise their predictions to a diverse array of proteins, leading to good performance in *most* regions at the expense of poor performance in unusual or poorly characterised regions, that do not occur often enough or contain enough clinically relevant variants to significantly reduce overall performance metrics. Further evidence of this is seen in the VEPs inability to identify several disease mechanisms that do not fit the standard loss-of-function paradigm among the most discordantly predicted variants and their general tendency to perform worse at gain-of-function and dominant-negative variants.

While the generation of MAVE datasets for all human proteins and isoforms is an aspirational goal of the Atlas of Variant Effects alliance (AVE), it is for the moment, not yet attainable (Fowler *et al*, 2023). The process of performing a mutational scan from conceptualisation to assay design, lab skills, troubleshooting, sequencing and post-processing is a highly skill-intensive process. Currently, relatively few groups are capable of performing the process end-to-end and generating high-quality datasets. So, for now, VEPs allow us to quickly and easily fill in the gaps with predictions that are still largely accurate.

In conclusion, while modern VEPs and MAVEs show a strong correlation, and tend to be in concordance for most clinically relevant variants, the cases where major discordance emerges sheds light on the weaknesses of both methods. By taking onboard the findings made here, there is scope for both MAVEs and VEPs to improve and better-reflect the true fitness of a greater proportion of variants.

## Methods

### Data acquisition and sources

Deep mutational scan datasets were obtained from several sources including their primary publication (Table 1), MaveDB (Esposito *et al*, 2019) and ProteinGym (Notin *et al*, 2023).

For the five variant effect predictors, AlphaMissense results were downloaded from their repository (https://zenodo.org/records/8208688). SaProt was run locally using instructions in their GitHub repository (https://github.com/westlake-repl/SaProt). ESM-1v was also run locally using their ‘zero shot variant effect prediction’ notebook as a template (https://github.com/facebookresearch/esm/tree/main/examples/variant-prediction). ESCOTT predictions were downloaded from the repository (https://zenodo.org/records/10670914). VARITY predictions were downloaded in bulk from the VARITY webserver (http://varity.varianteffect.org/).

Pathogenic and benign variants were labelled using ClinVar (06/08/2024 release). Putatively benign variants were labelled using gnomAD version 4.1.

Protein feature annotations were obtained from UniProt (https://www.uniprot.org/) and from InterPro (https://www.ebi.ac.uk/interpro/).

AlphaFold predicted local distance difference test (pLDDT) scores were extracted from the AlphaFold PDB-format files (https://alphafold.ebi.ac.uk/).

Relative solvent exposed surface area (RSA) was calculated from first biological assembly files in the protein databank (PDB) (https://www.rcsb.org/).

### ROC AUC calculation

Area under the receiver operating characteristic curve (AUROC) was calculated using the roc_auc_score() function of the sklearn python package (0.18.1). Prior to AUROC calculation, any dataset with an inverted scale was deducted from 1, to ensure that a score of 0 represented wild type-like variants and 1 represented the greatest fitness effects.

For the generation of Figure 1, the pathogenic dataset was obtained from ClinVar, filtered for pathogenic and likely pathogenic variants with a 1-star review status or higher. The putatively benign dataset composed of gnomAD variants (passing the internal filter). The AUROC calculation was bootstrapped 1000 times with independent selection with replacement from the pathogenic and putatively benign datasets.

### Rank normalisation

To perform rank normalisation, any dataset with an inverted scale was deducted from 1, to ensure that a score of 0 represented wild type-like variants and 1 represented the greatest fitness effects. Scores were converted into ranks using the dataframe.rank() function of the pandas python package (1.1.5). The default behaviour of the rank function was used, which assigns the average rank of the group to any tied values. Following ranking, the ranked data was normalised min-max normalised between values of 0 and 1.

Where multiple columns of data are compared to each other, all values missing from at least one column were removed from all columns prior to ranking to ensure that ranks were all calculated with the same variants.

### Interface classification

The classifications of surface, interior and interface were assigned as previously described (Levy, 2010) by RSA. RSA was calculated from the first biological assembly of each protein in the PDB. Interfaces are defined as areas where the RSA changes between the monomeric and complex assembly structures. Surface residues have an RSA of 25% or higher while interior residues have an RSA less than 25%. Among interface residues, support residues have an RSA of less than 25% in both the monomer and complex. Interface rim residues have an RSA of 25% or higher in both the monomer and complex. Core interface residues have an RSA of 25% or higher in the monomer but less than 25% in the complex.

We previously found that interface rim residues were very similar to surface residues in terms of properties and pathogenic variant enrichment, so all calculations using these definitions include the interface rim as surface resides (Livesey & Marsh, 2022).

## Acknowledgements

This project was supported by core funding from the Medical Research Council to the MRC Human Genetics Unit (MC_UU_00035/9).

## Supplementary Figures

**Figure S1.**
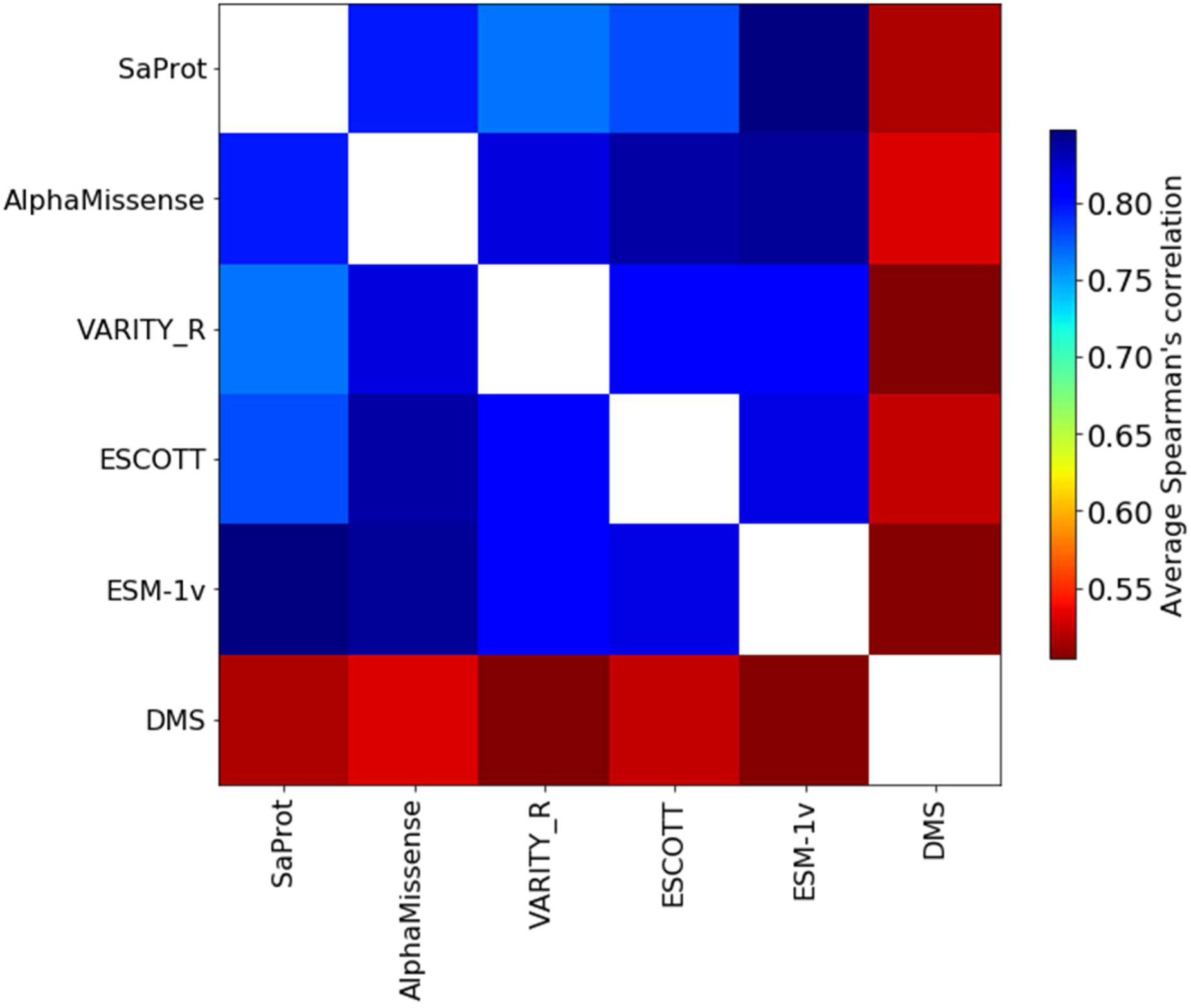
Average Spearman’s correlation between each of the VEPs and the MAVE data across all 37 proteins. Spearman’s correlation in each pairwise comparison is averaged across each of the 37 MAVE datasets.

**Figure S2.**
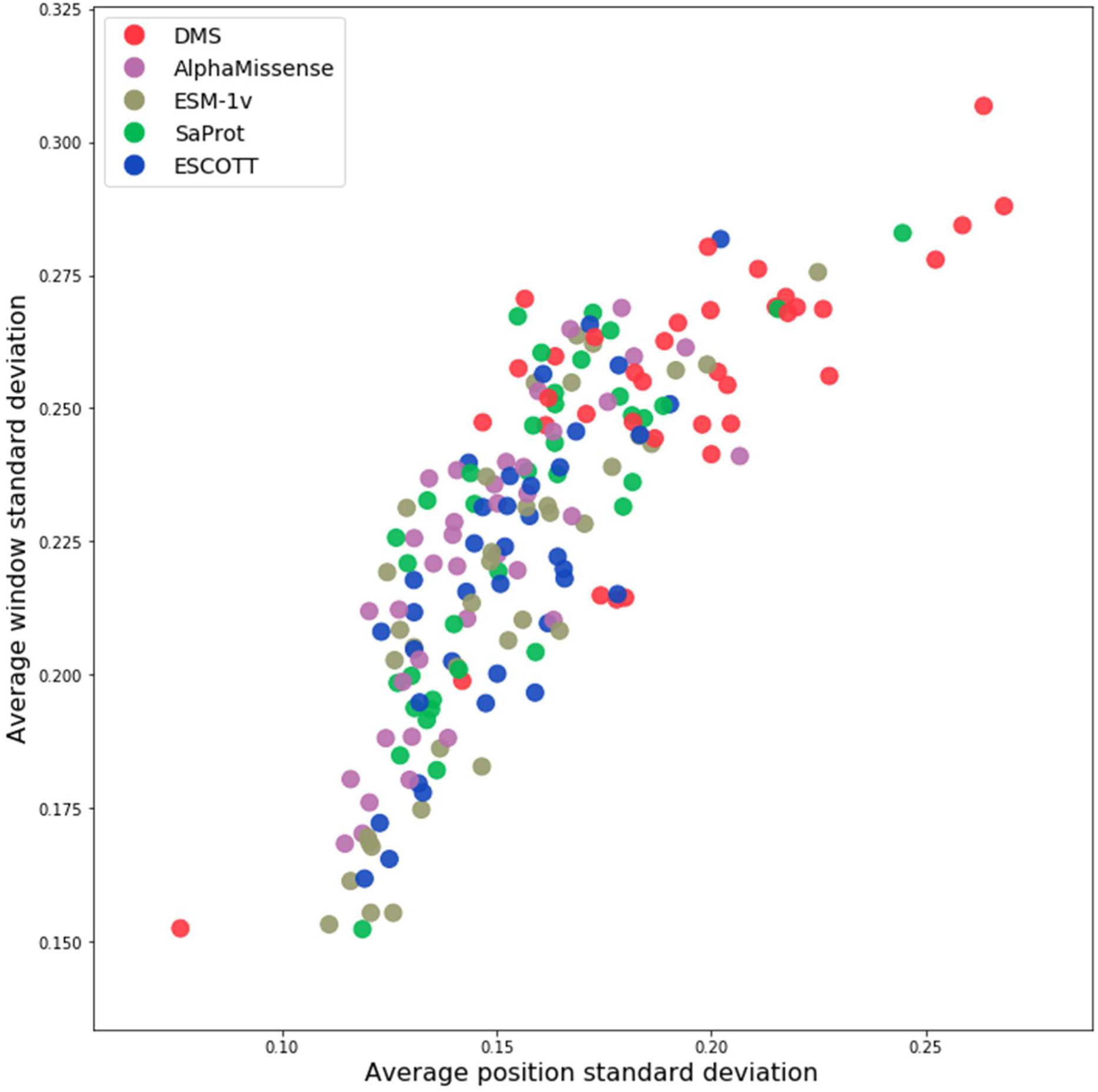
Average standard deviation of MAVE and VEP score sets. The average standard deviation of the normalised rank scores of all possible amino acid substitutions at each position in a protein plotted against the average standard deviation of all possible amino acid substitutions in 10 amino acid bins for each protein.

**Figure S3.**
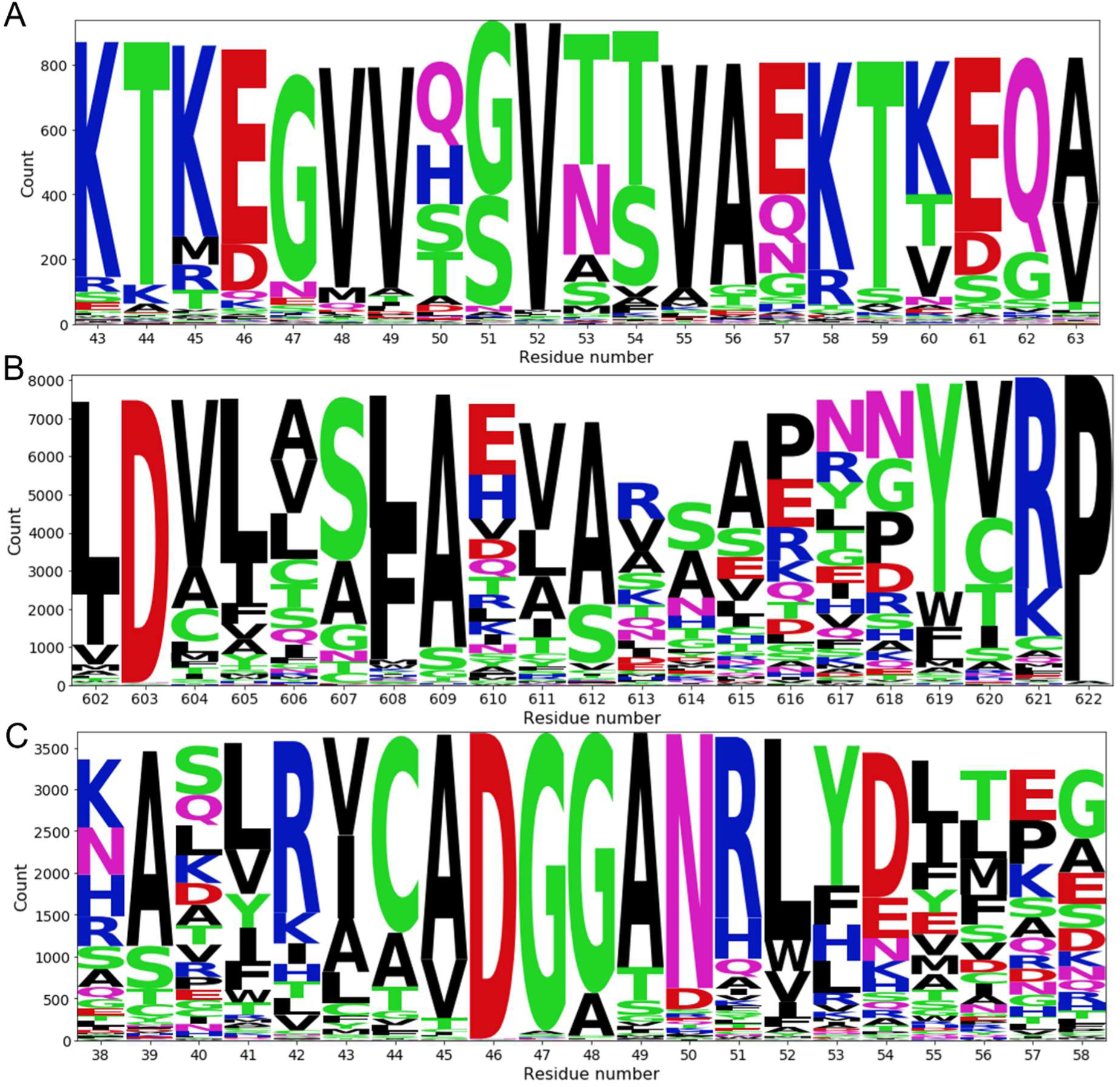
Logo plots representing the number of each amino acid present at each position in a multiple sequence alignment of three proteins. Multiple sequence alignments were generated by MMseqs2, the size of each letter represents the total number of that particular amino acid at each aligned position. The differences in column size are due to the removal of gap characters and non-standard amino acid notations from the analysis. **A.** A representation of the α-synuclein alignment around the location of pathogenic variant A53T. **B.** A representation of the *MSH2* alignment around the location of allegedly benign variant S612P. **C.** A representation of the *TPK1* alignment around the location of pathogenic variant G48V.

