## Supplementary material for "Why variant effect predictors and multiplexed assays agree and disagree": File SF1

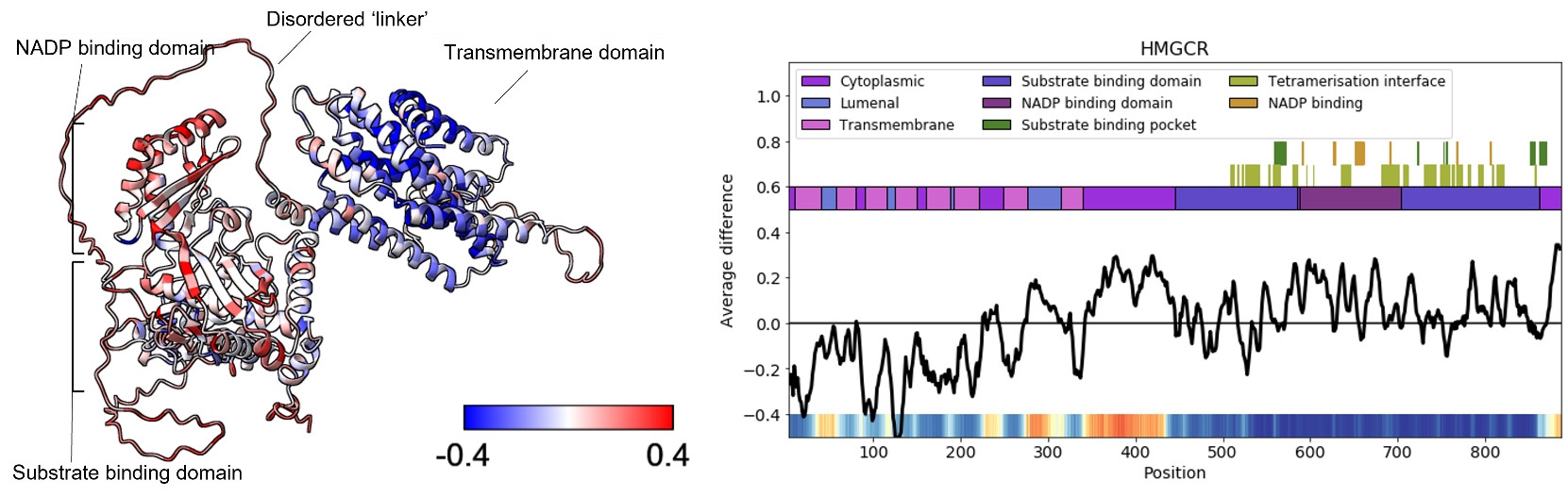
**Supplemental file 1.** Mean normalised difference between MAVEs and VEPs plotted across the protein structure and sequence for 37 proteins. Each protein structure was obtained from AlphaFold2, sequence annotations were obtained from UniProt and the conserved domains database. Positive average difference indicates regions where the MAVE predicts variants as more pathogenic while negative average differences indicate regions where the VEPs consider variants more pathogenic.

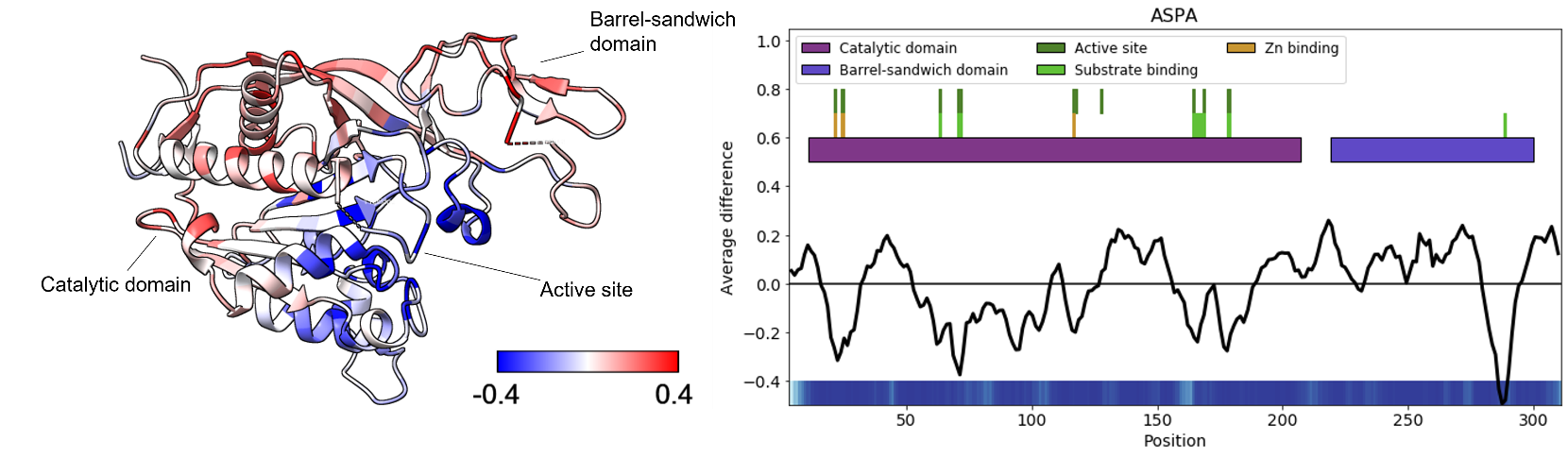

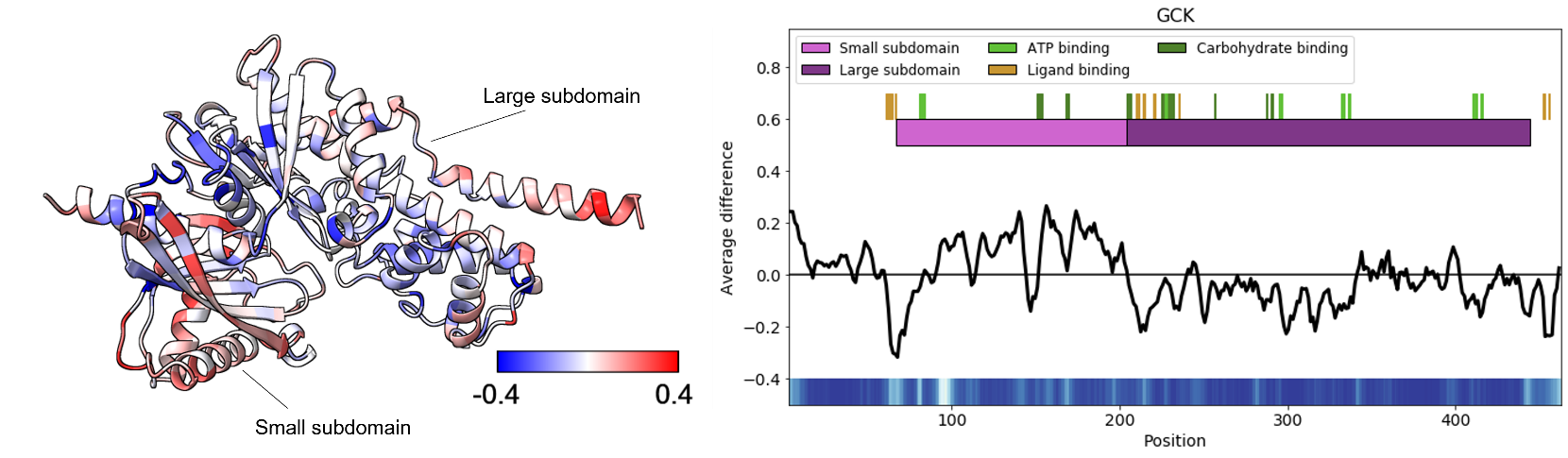

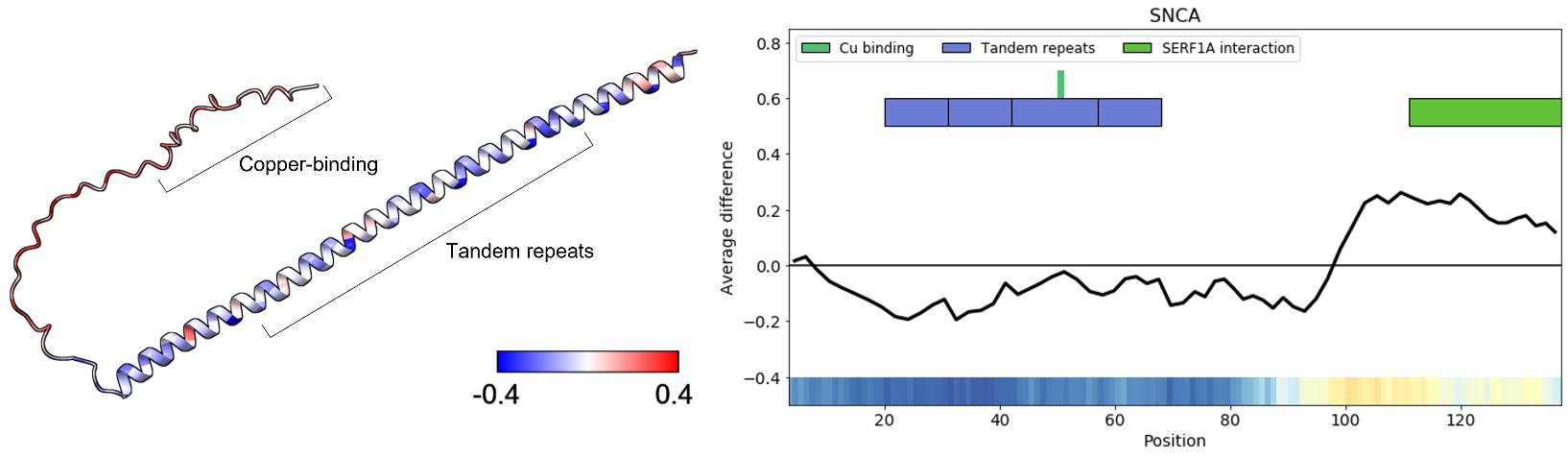

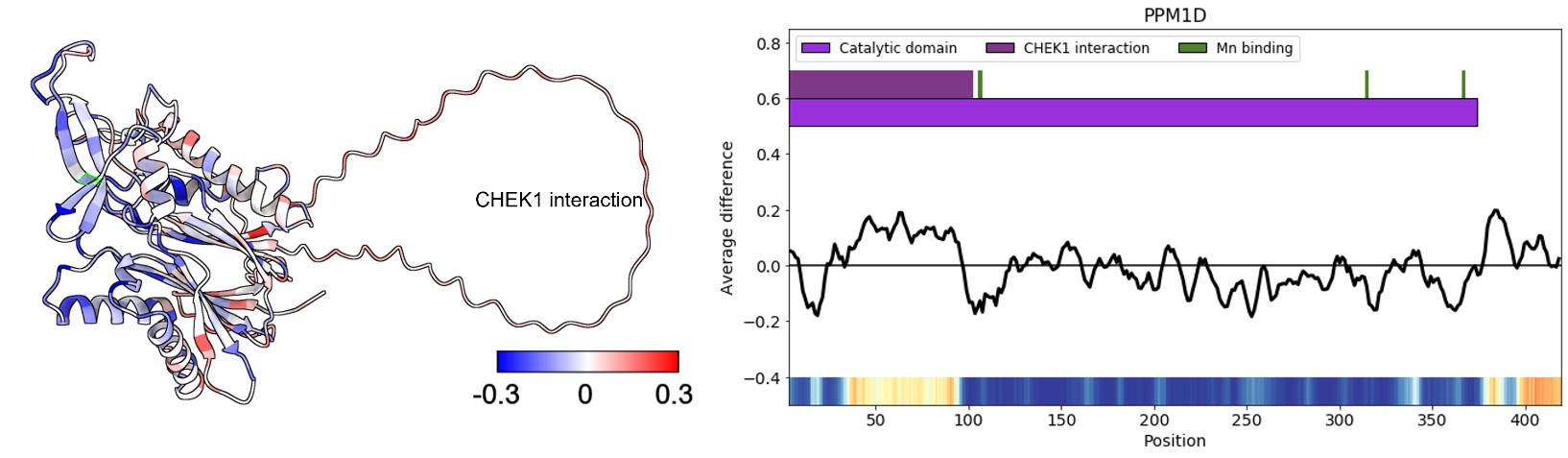

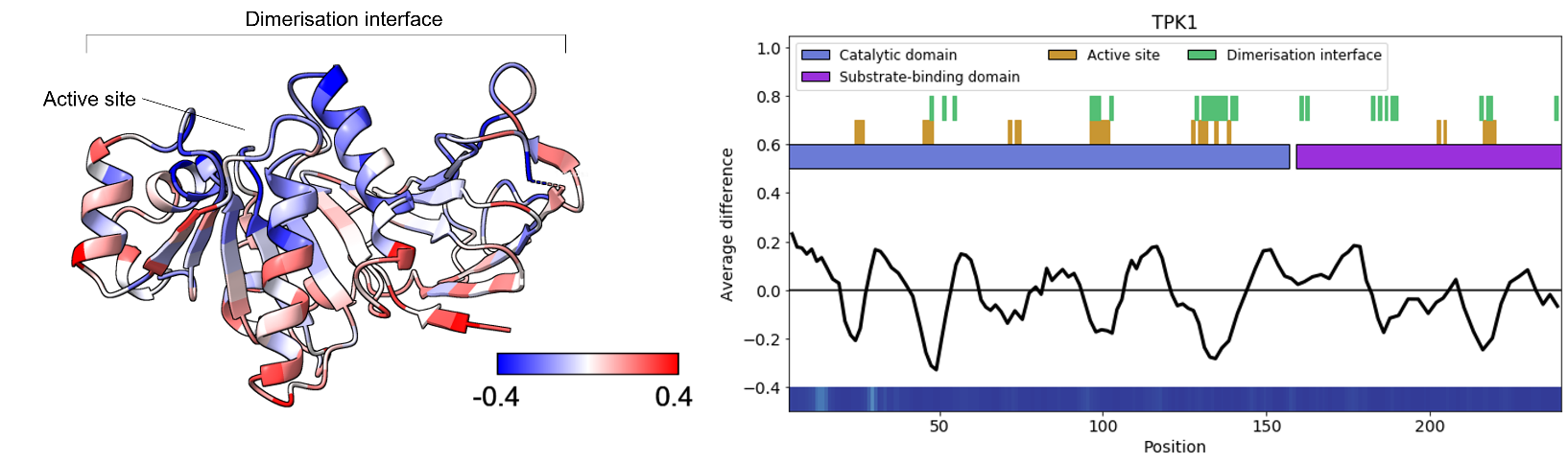

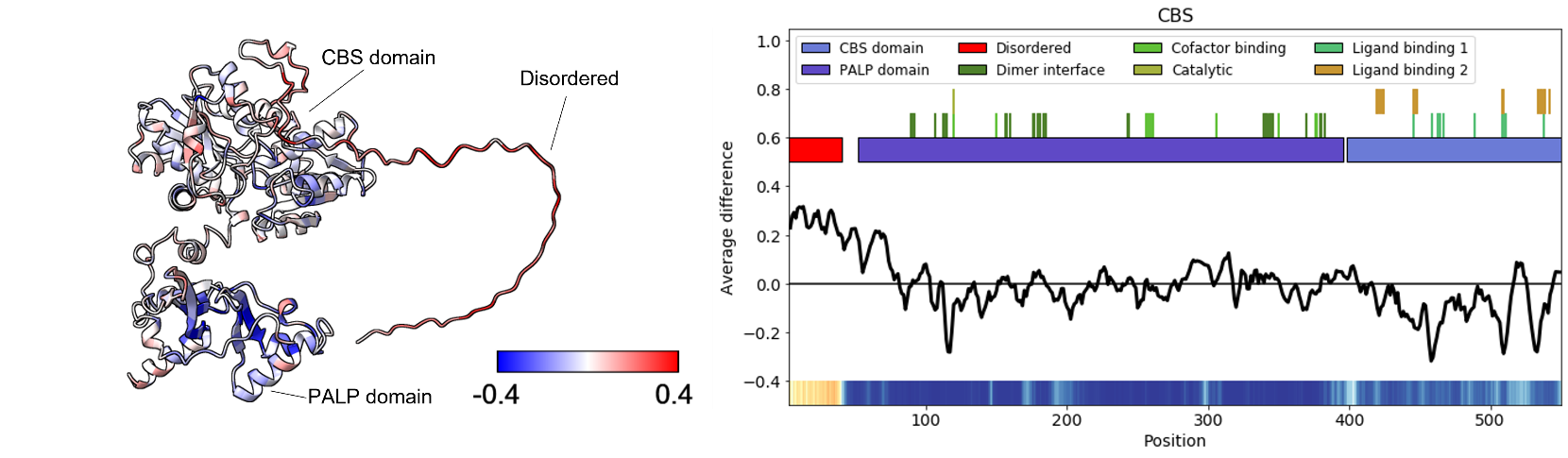

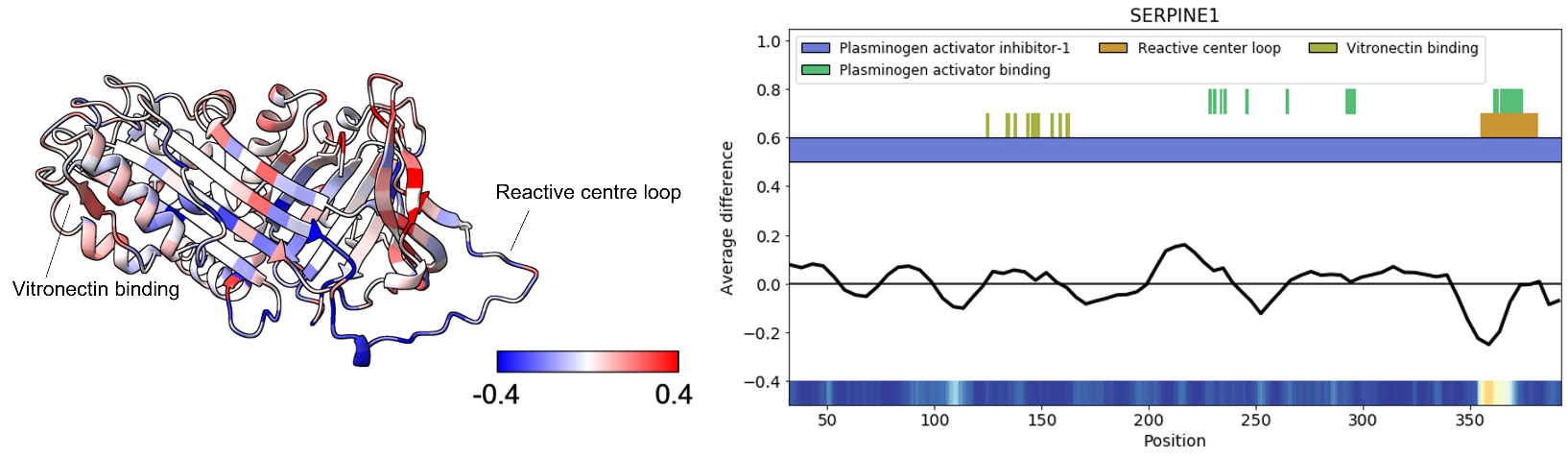

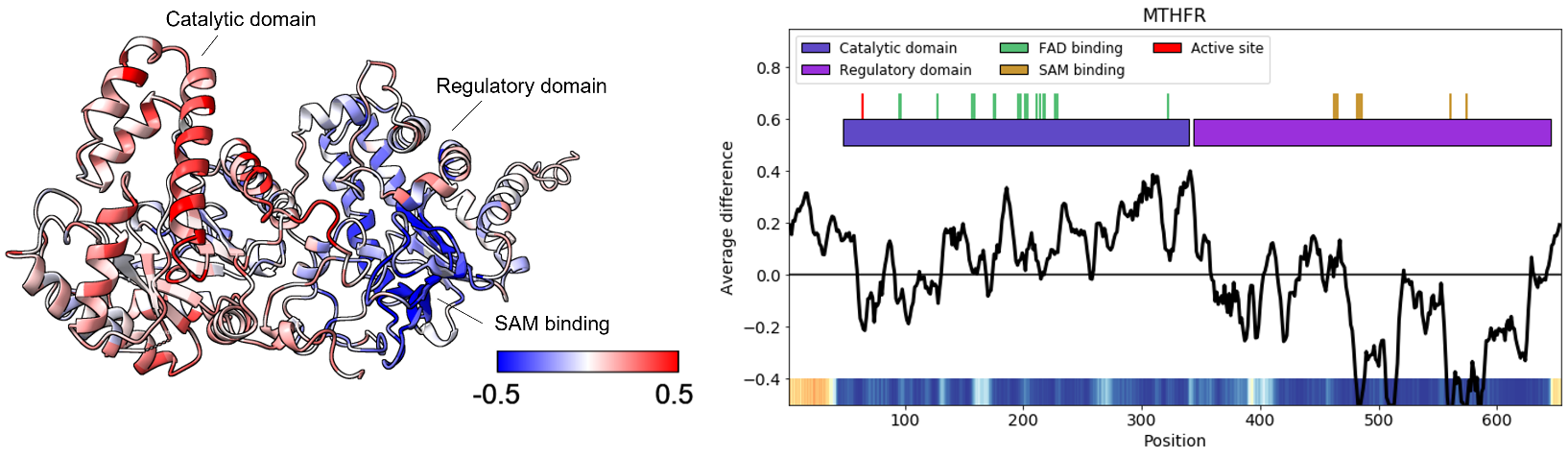

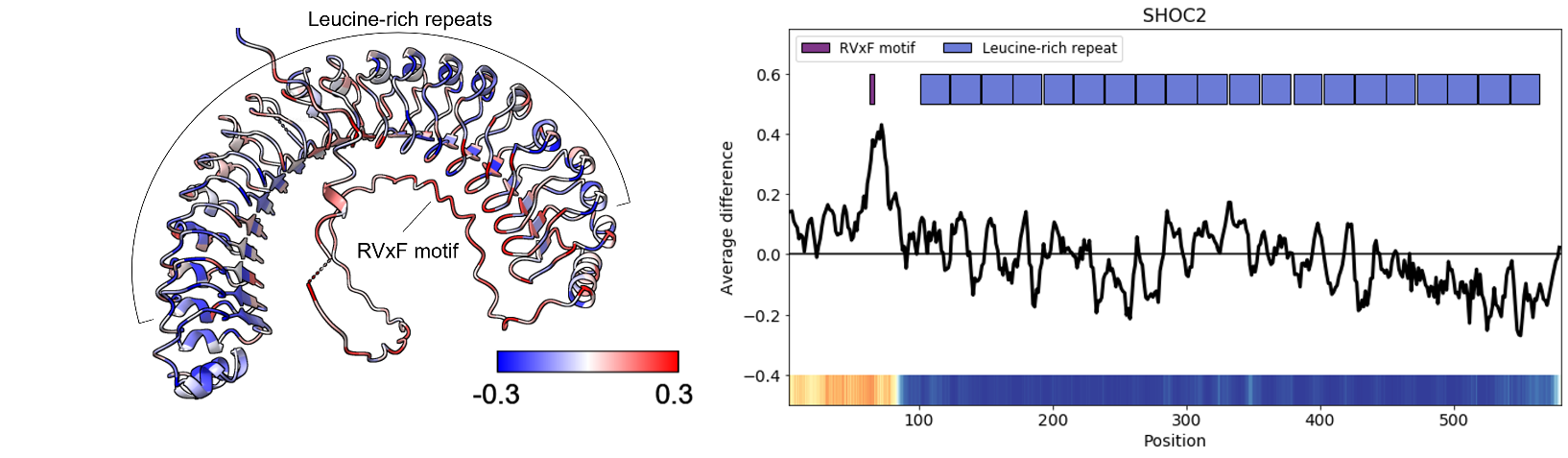

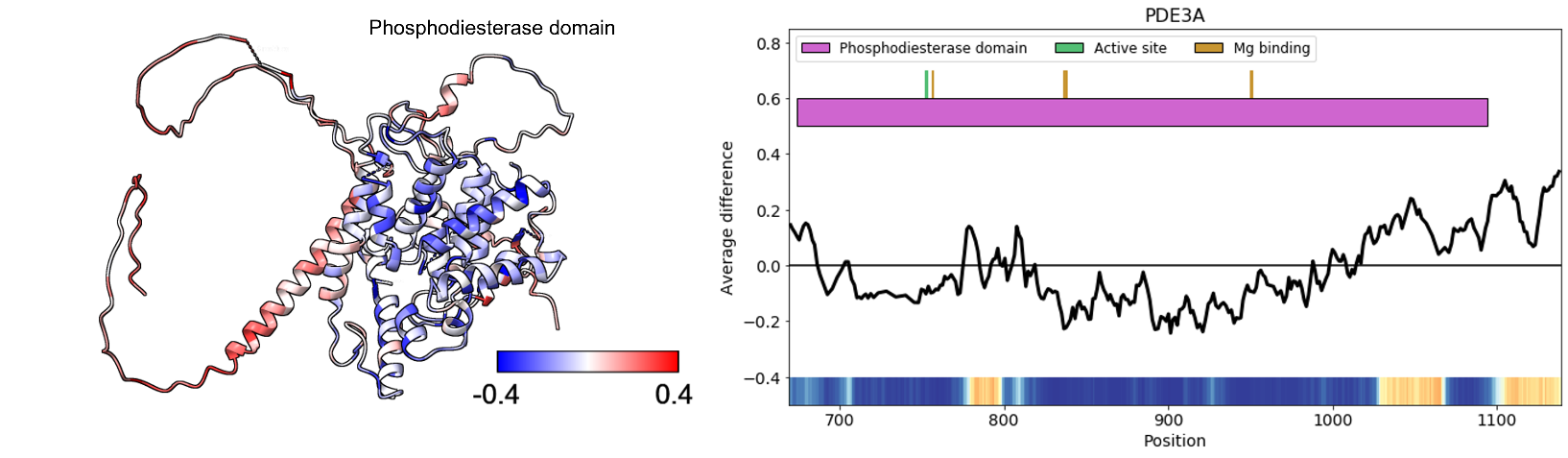

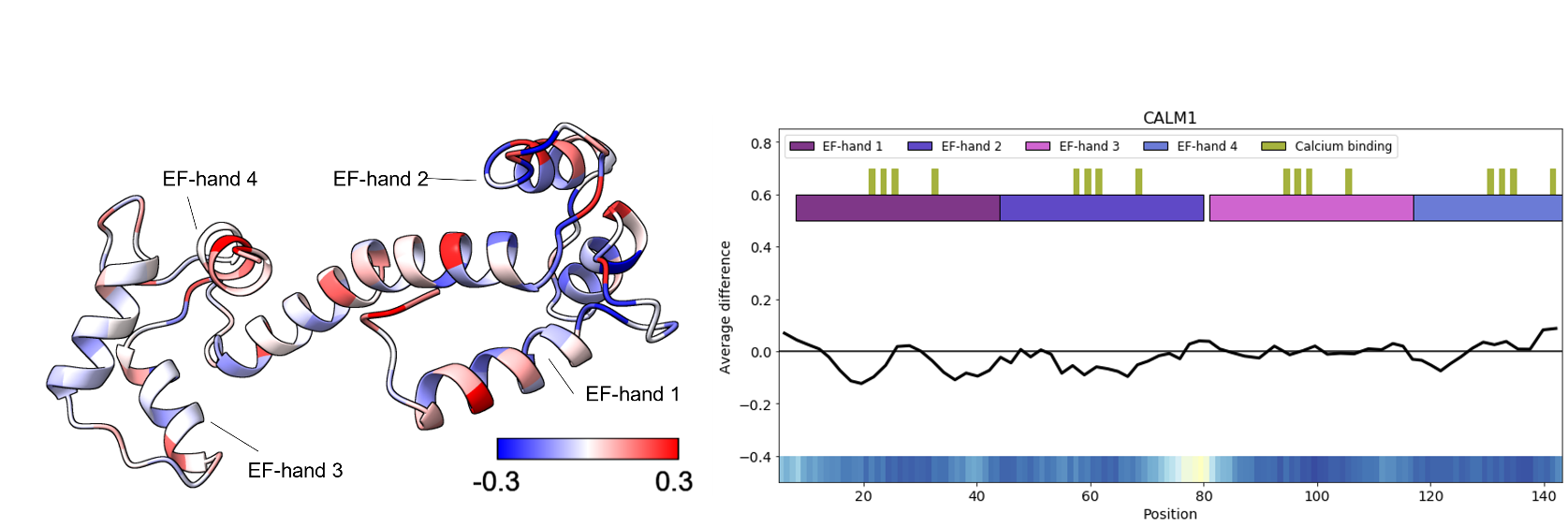

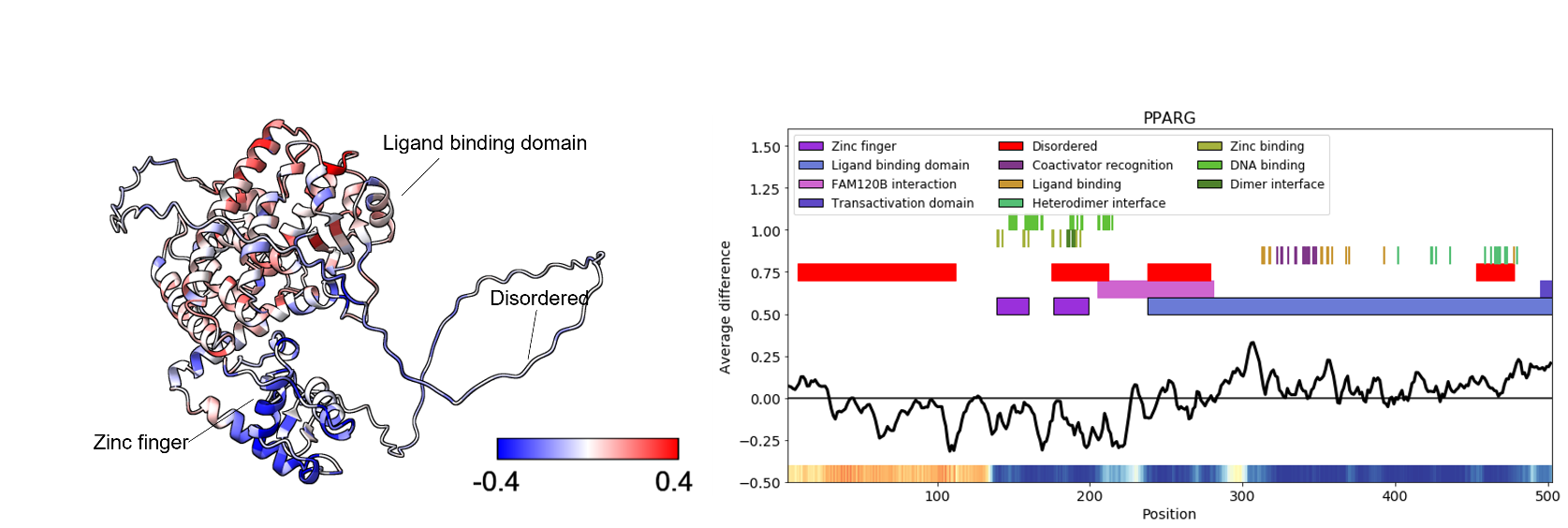

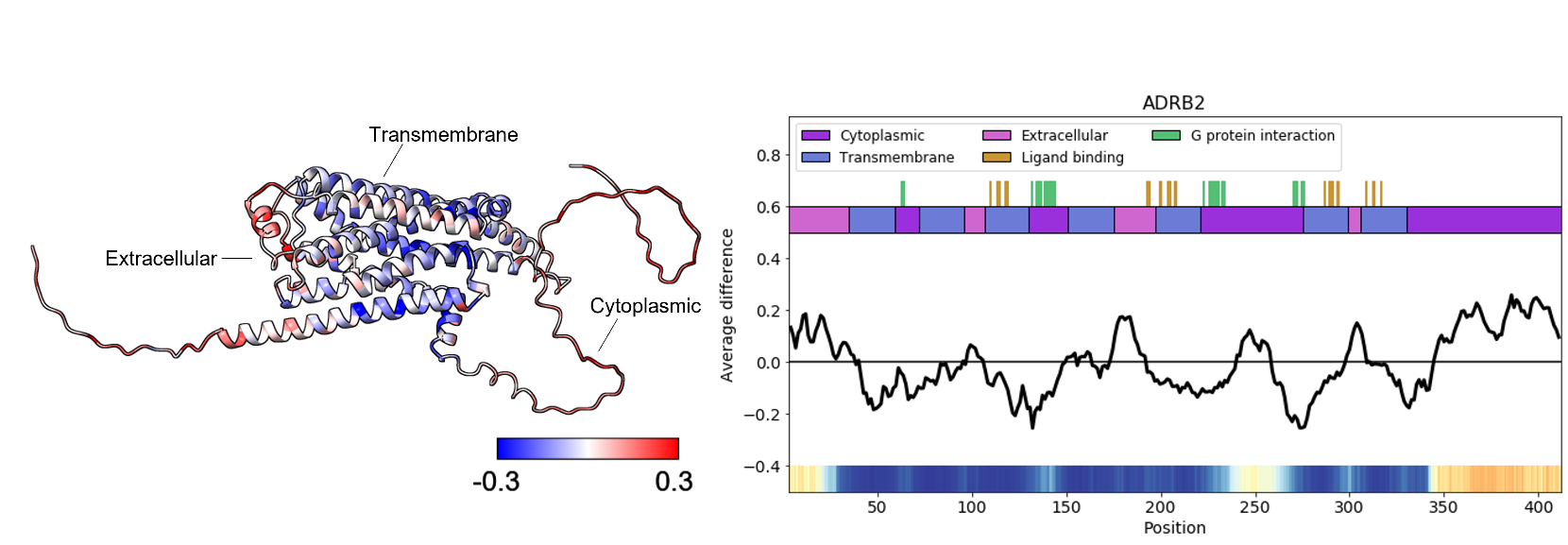

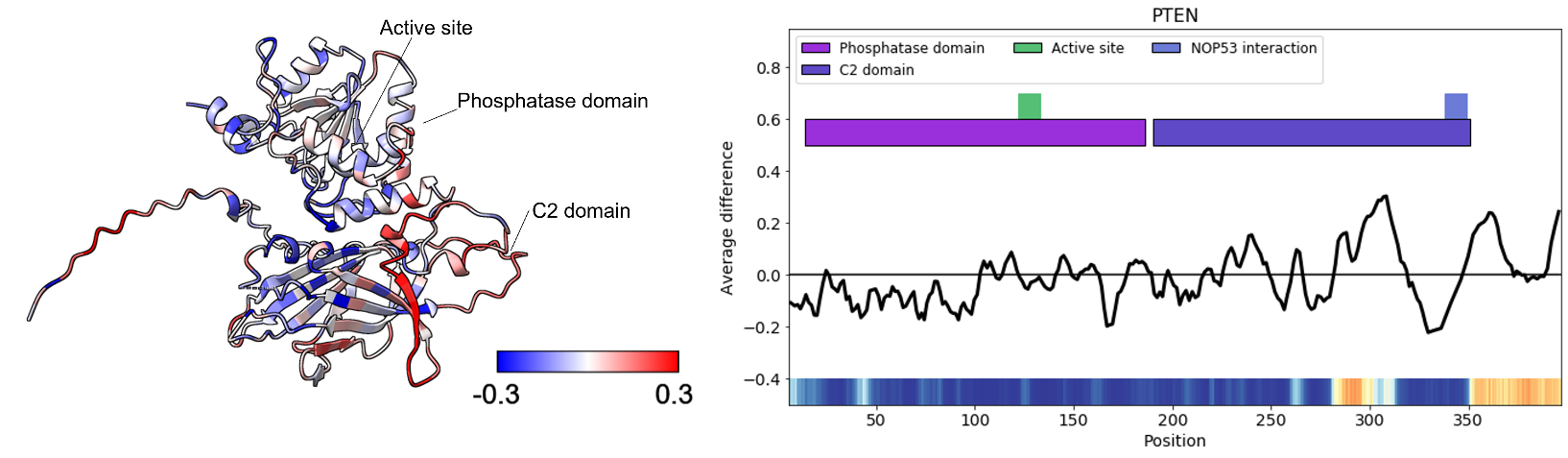

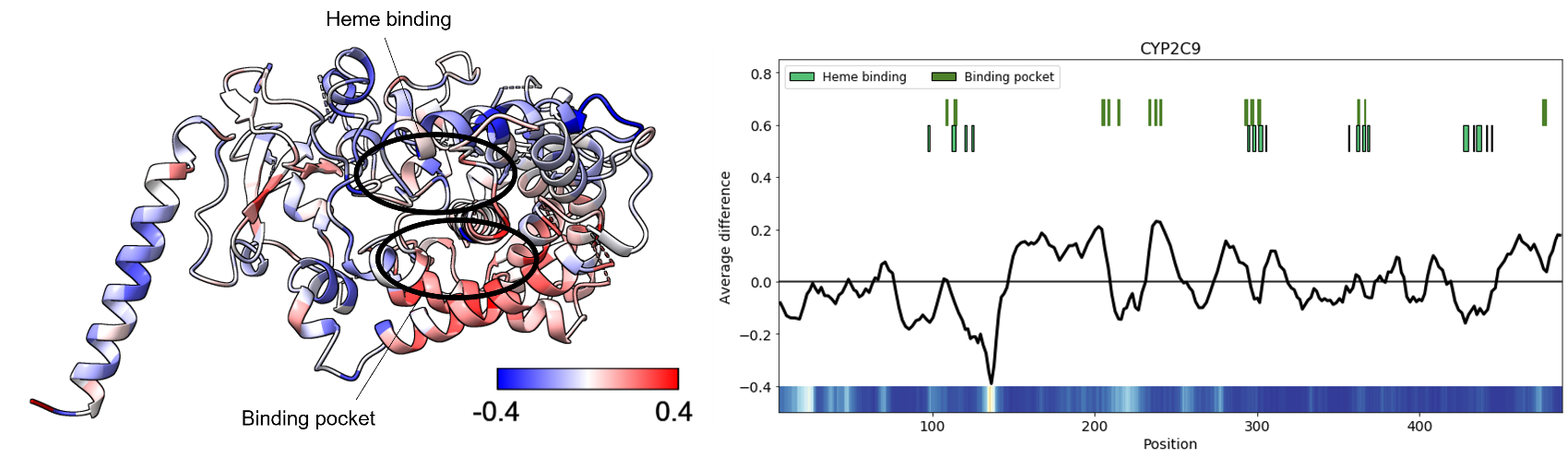

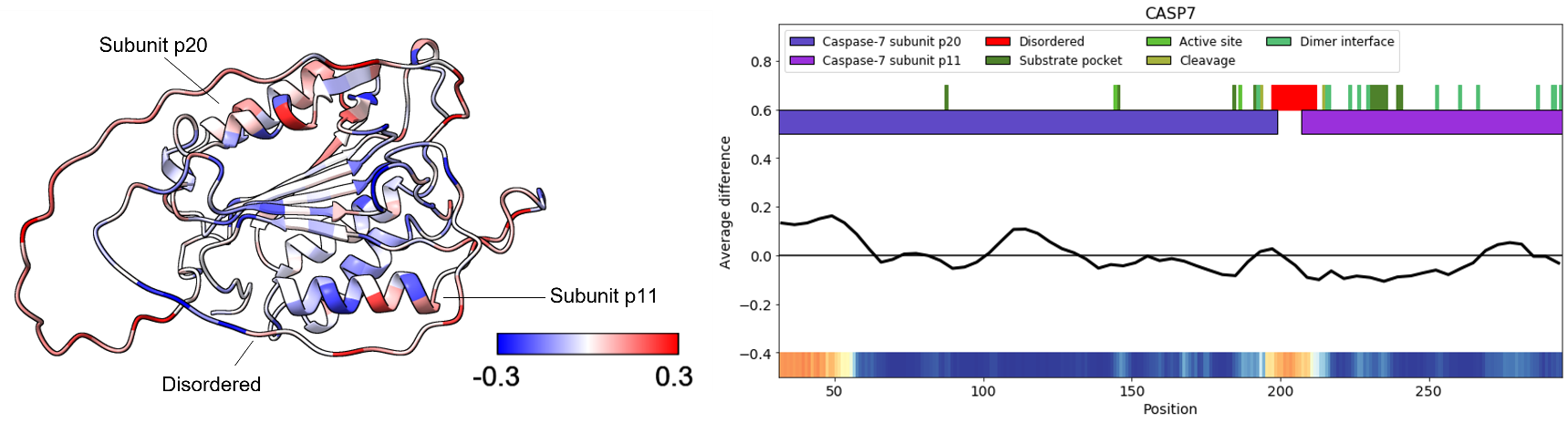

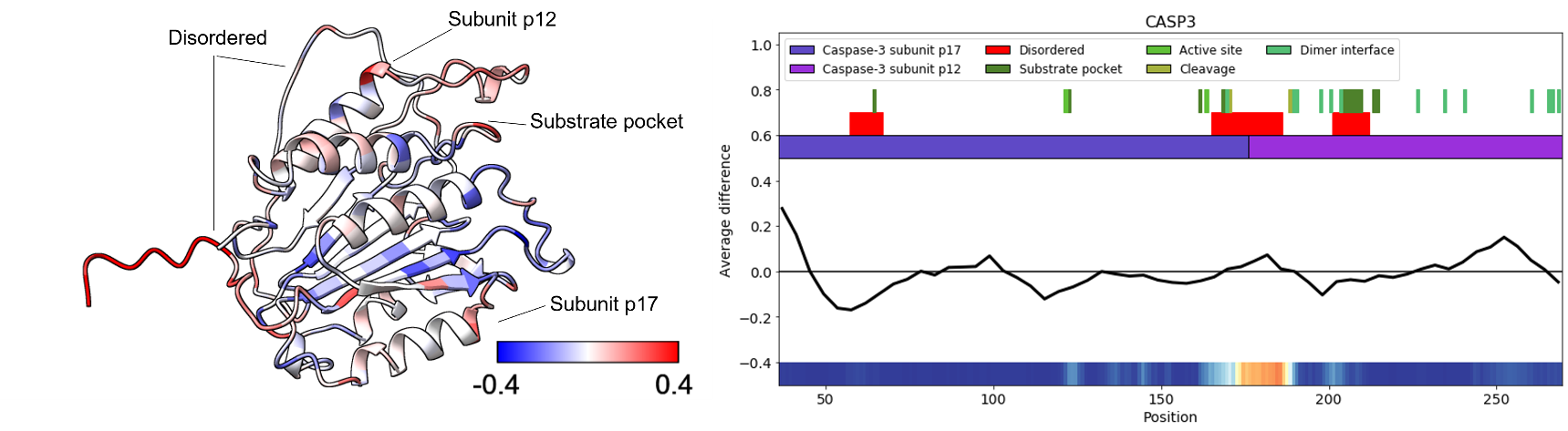

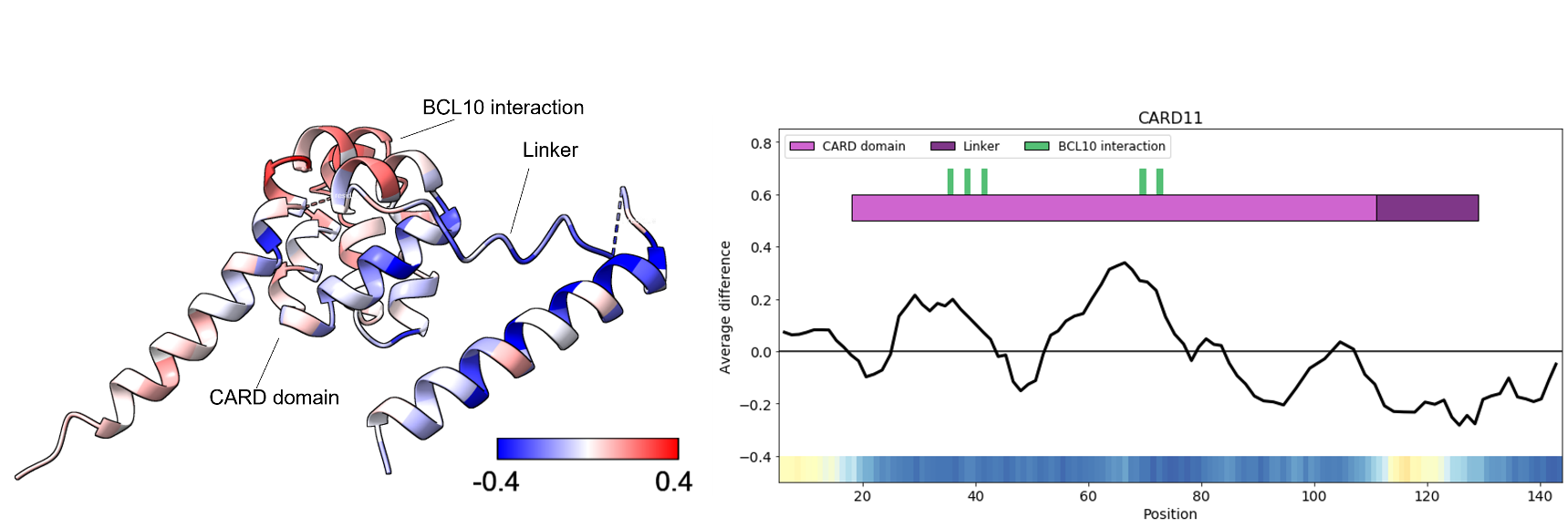

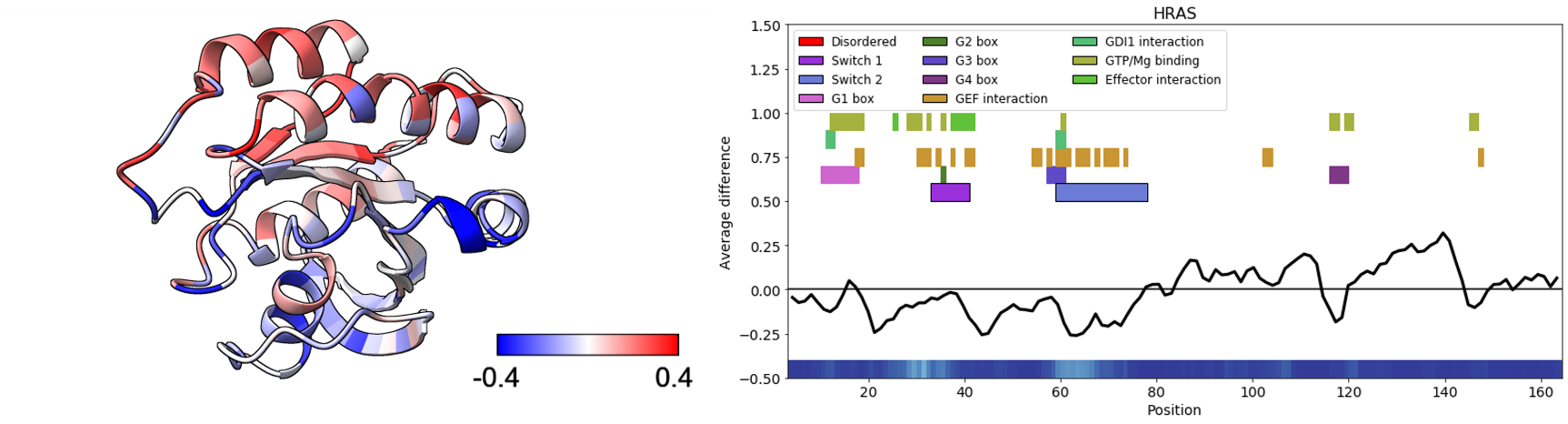

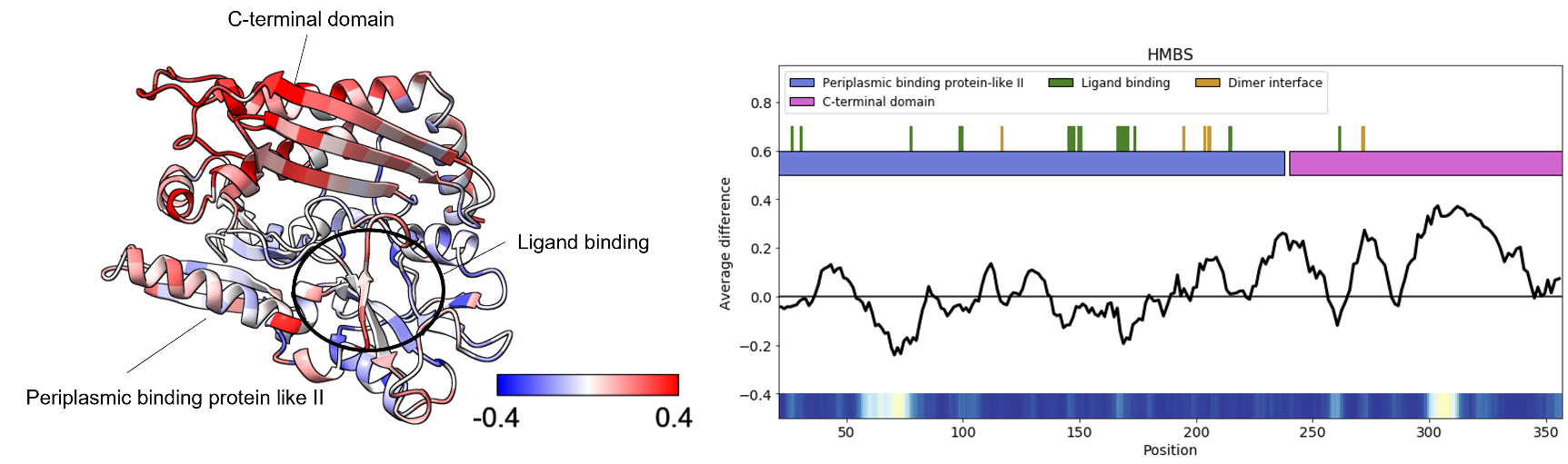

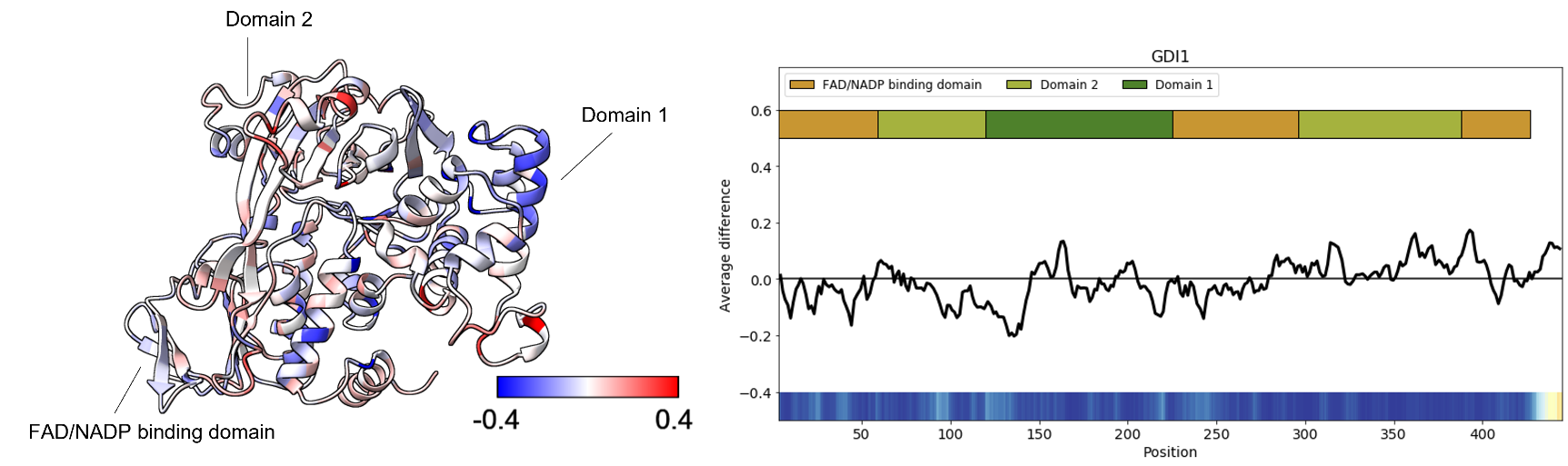

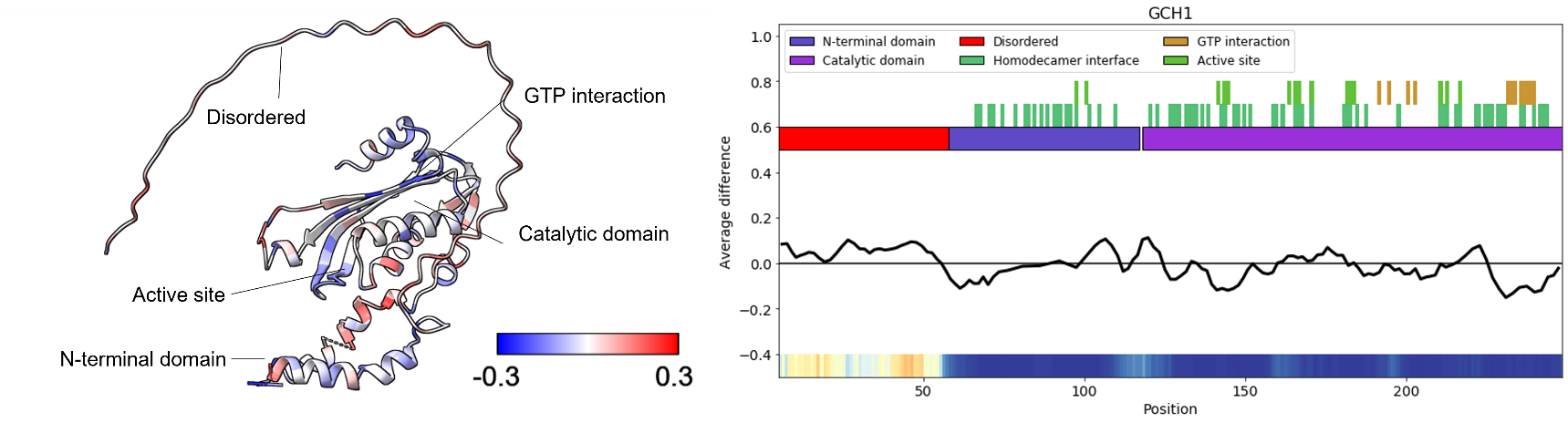

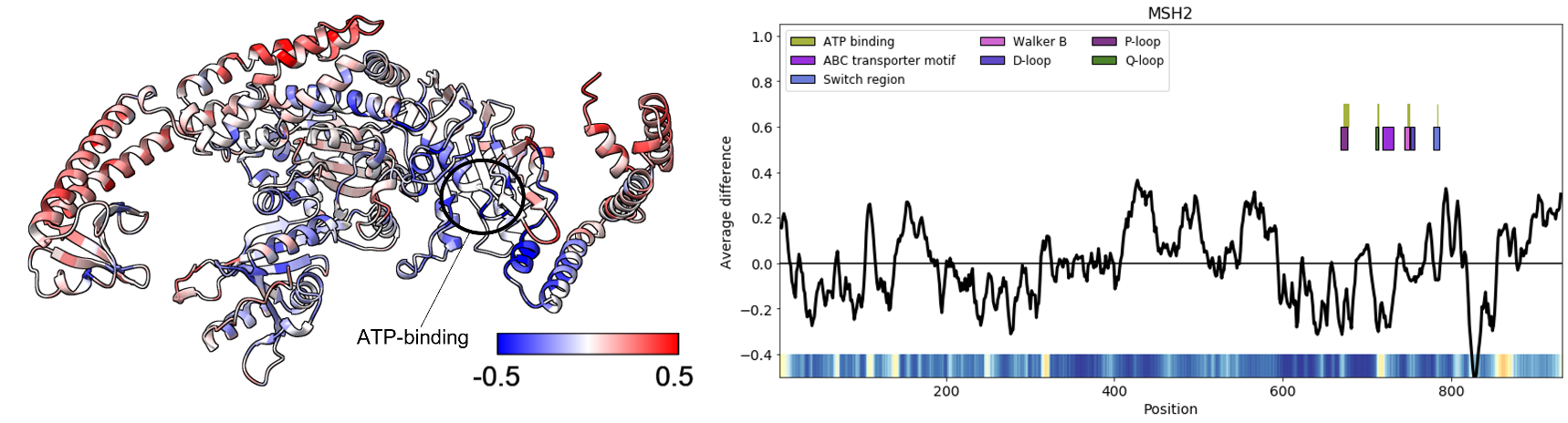

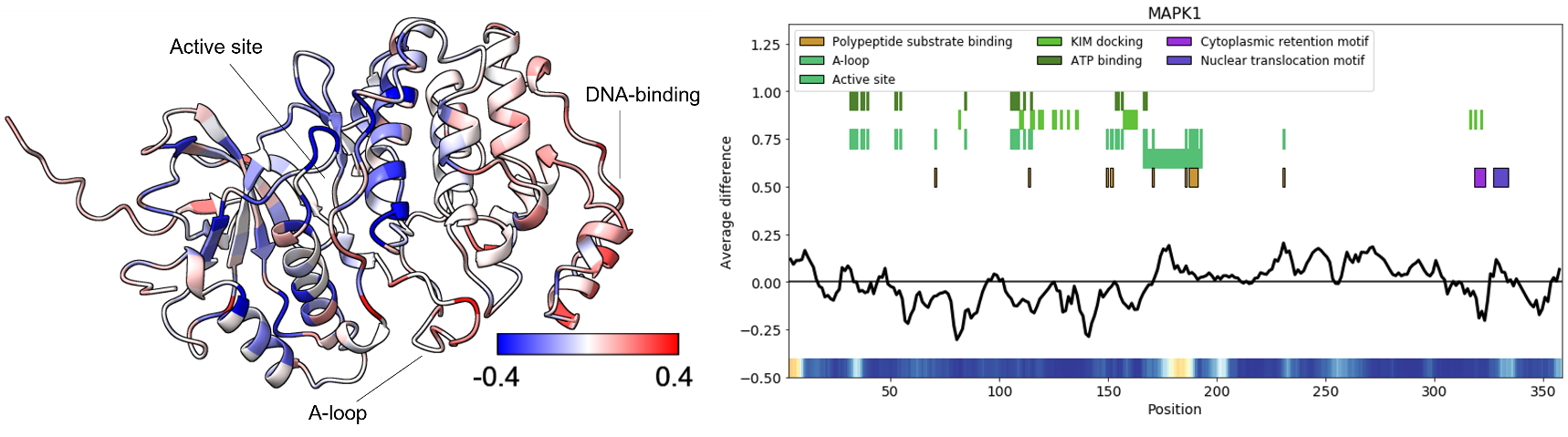

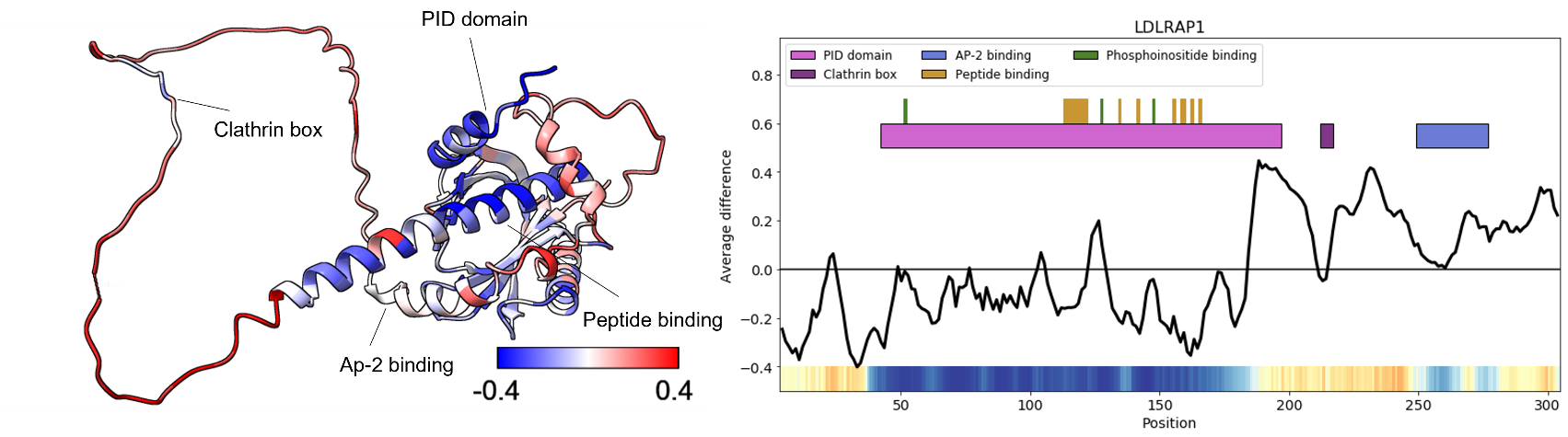

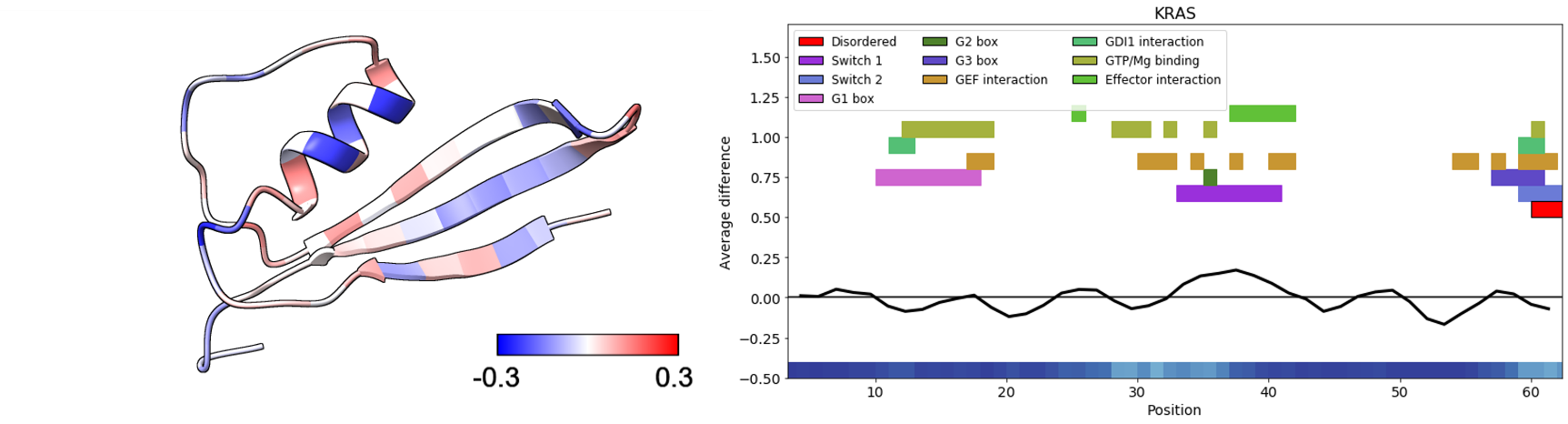

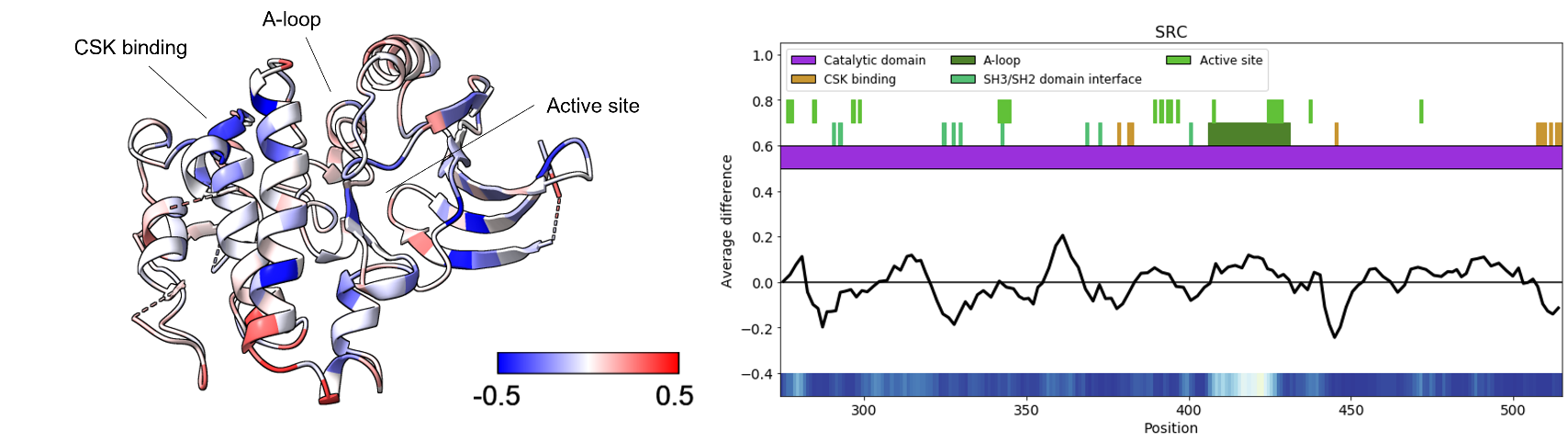

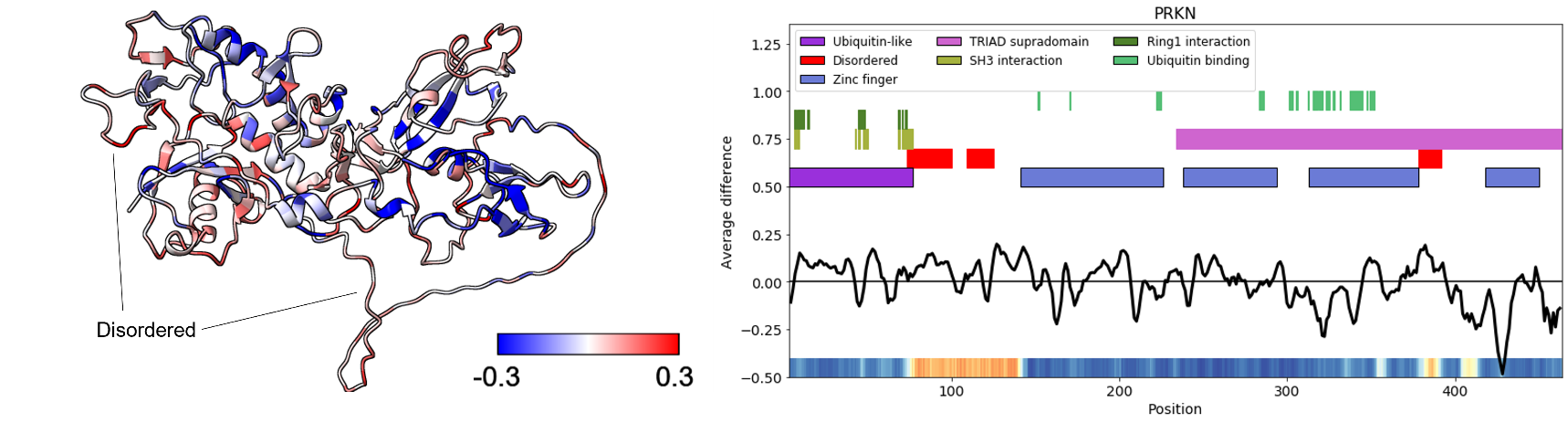

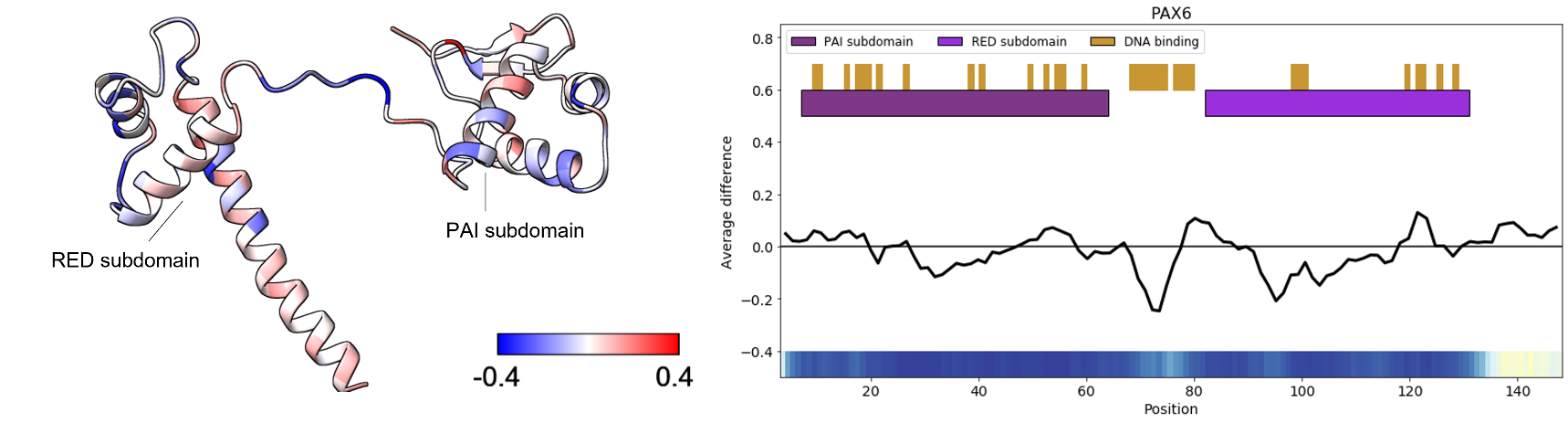

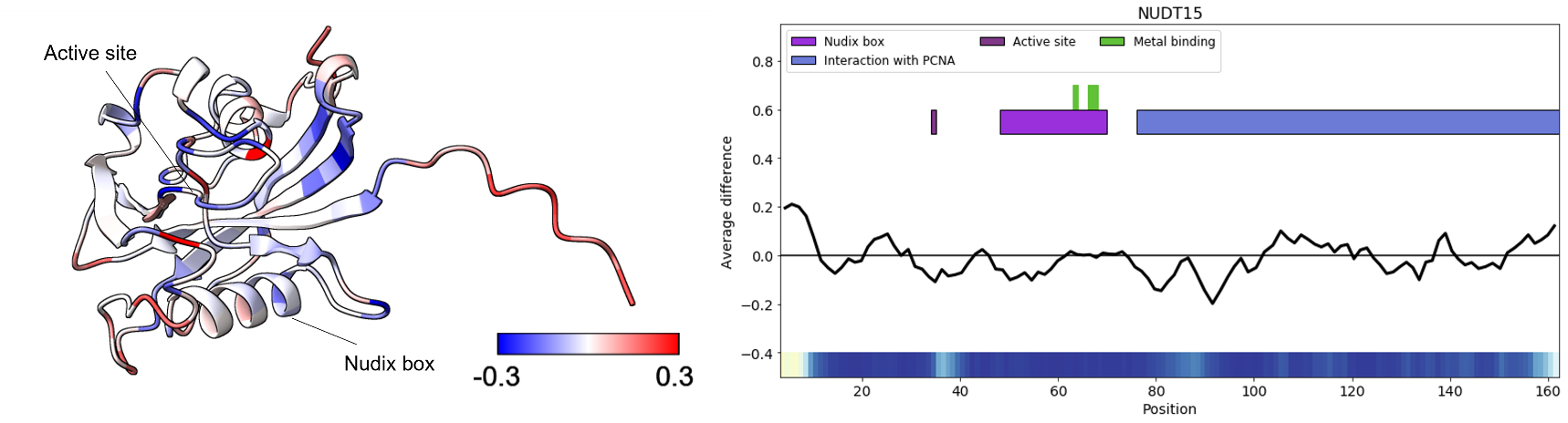

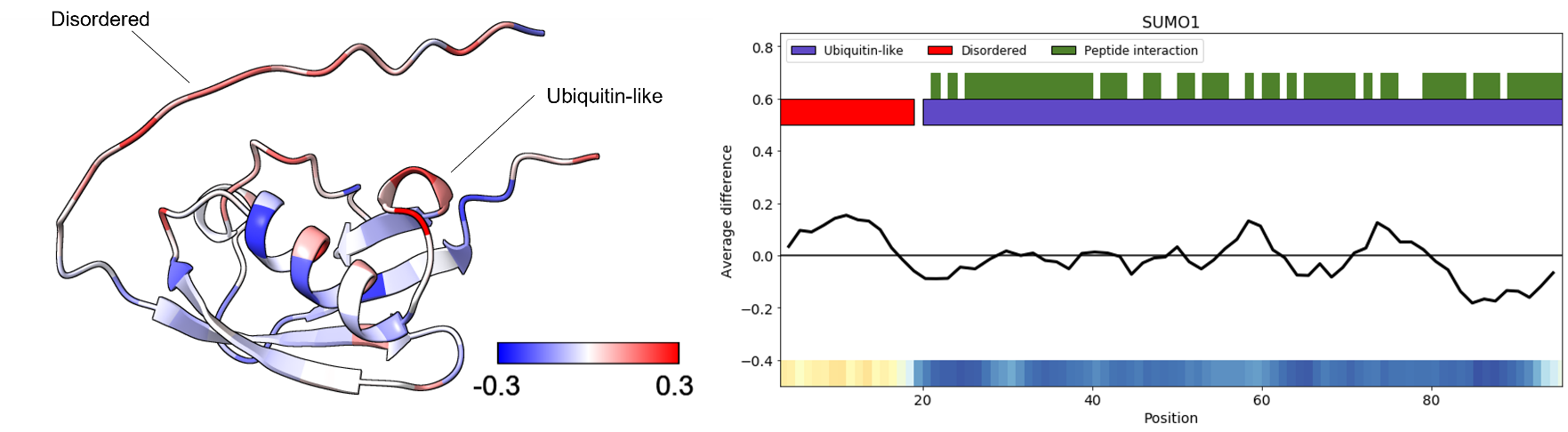

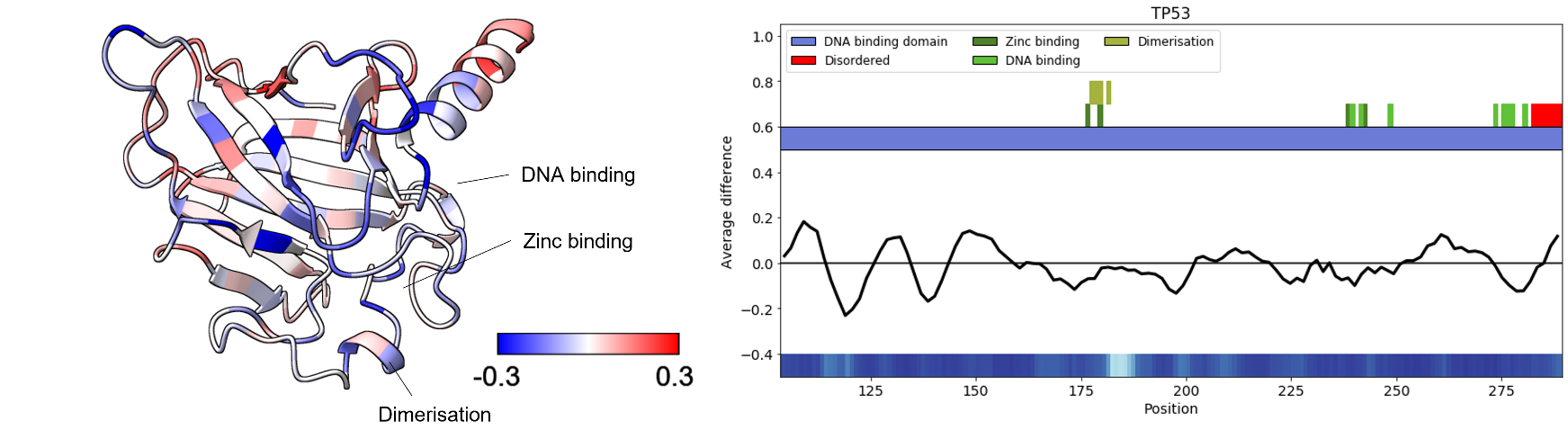

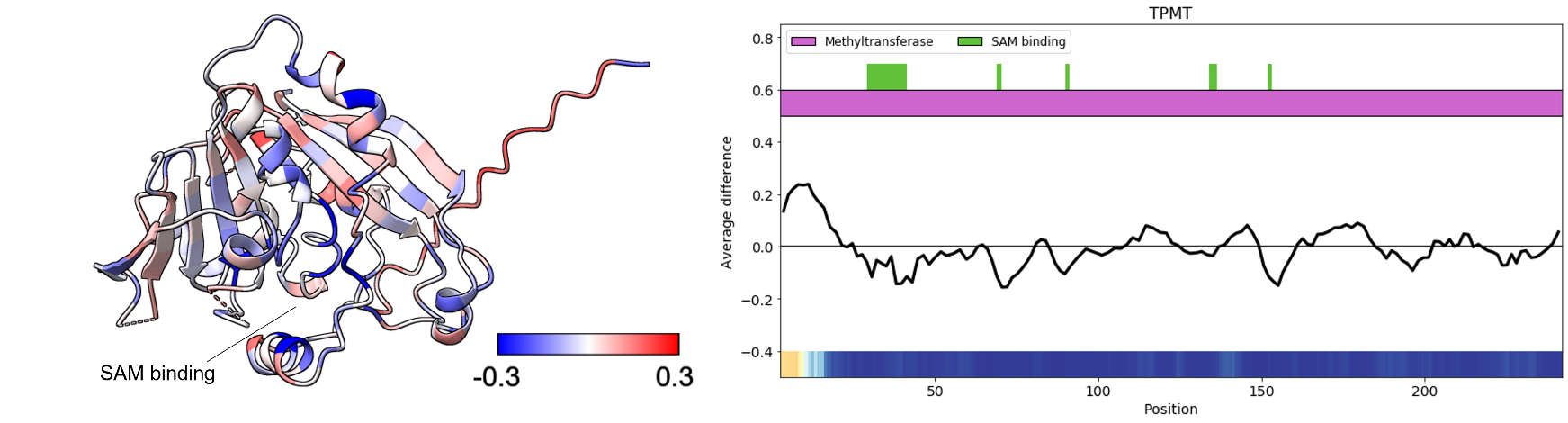

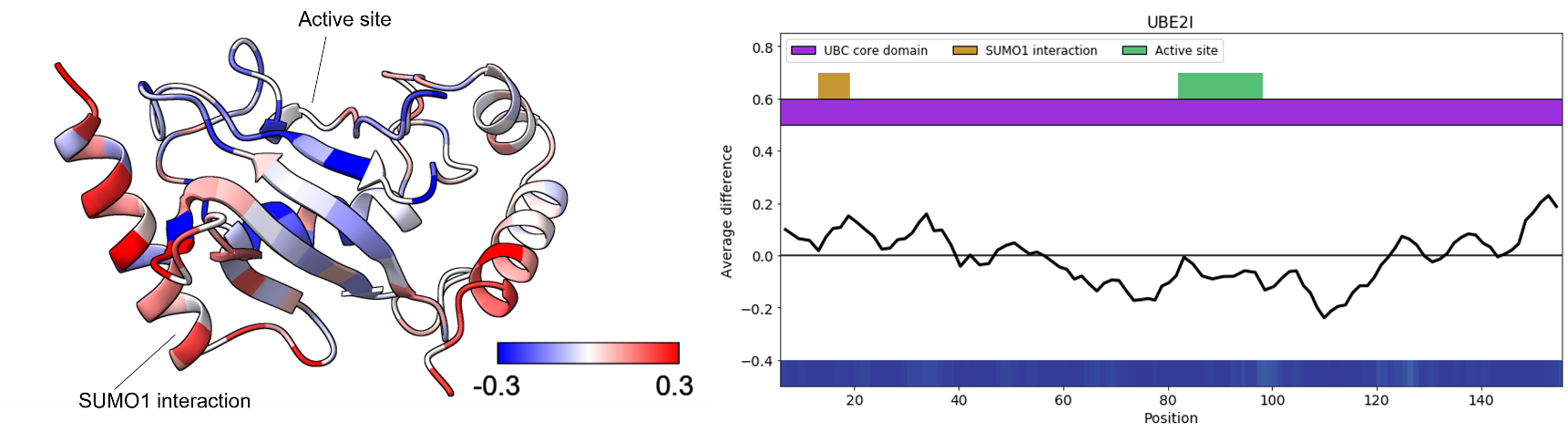

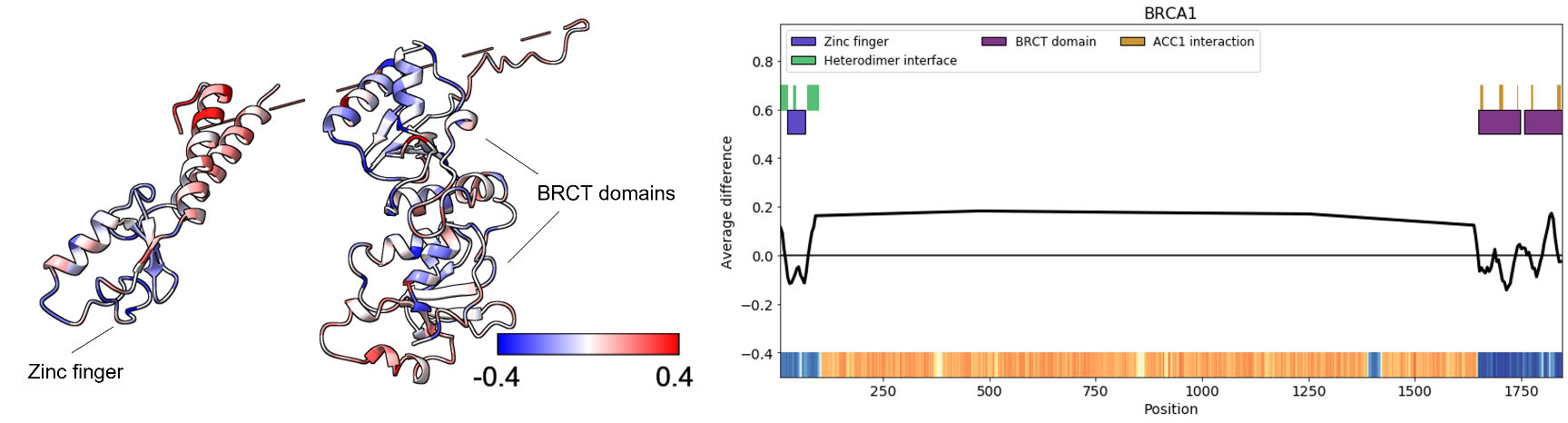

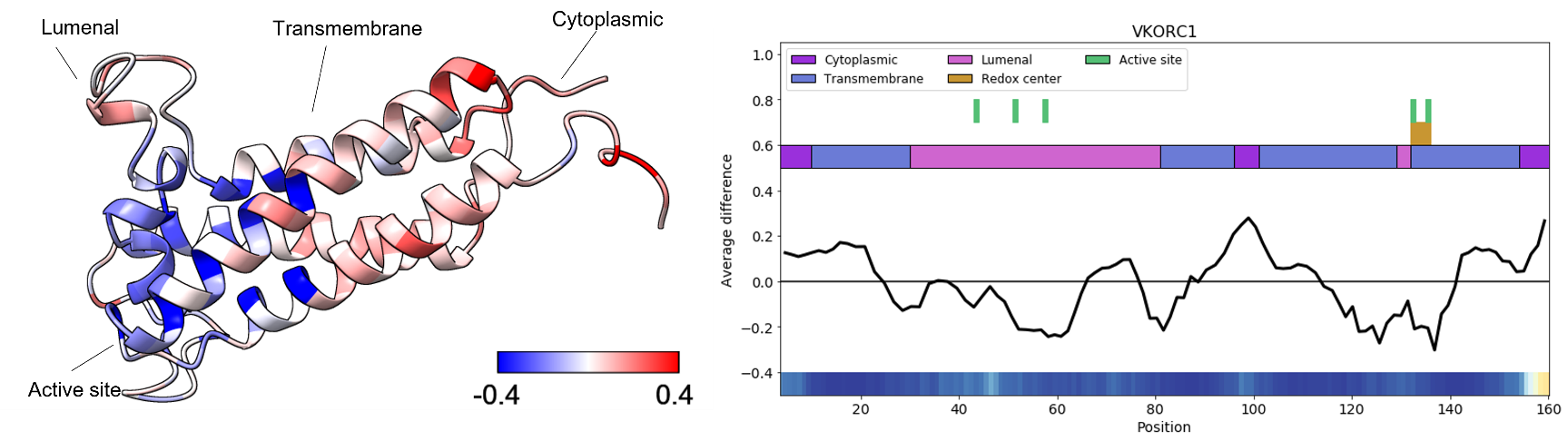
